# MALDI-TIMS-MS^2^ Imaging and Annotation of Natural Products in Fungal-Bacterial Co-Culture

**DOI:** 10.1101/2025.05.11.653367

**Authors:** Robert A. Shepherd, Gordon T. Luu, Laura M. Sanchez

## Abstract

Mass spectrometry imaging (MSI) is a powerful tool for monitoring the spatial distributions of microbial metabolites directly from culture. MSI can identify secretion and retention patterns for microbial metabolites, allowing for the assessment of chemical communication within complex microbial communities. Microbial imaging via matrix-assisted laser desorption/ionization (MALDI) MSI remains challenging due to high sample complexity and heterogeneity associated with the required sample preparation, making annotation of molecules by MS^1^ alone challenging. The implementation of trapped ion mobility spectrometry (TIMS) has increased the dimensionality of MALDI-MSI experiments, allowing for the resolution of isomers and isobars, and can increase sensitivity of metabolite detection within complex samples. Parallel reaction monitoring – parallel accumulation serial fragmentation (prm-PASEF) leverages TIMS to enhance the targeted acquisition of MS^2^ data by increasing the number of precursors that can be fragmented in a single acquisition. Recently, imaging prm-PASEF (iprm-PASEF) has been developed to provide more accurate annotation from MALDI-TIMS-MSI datasets through the inclusion of MS^2^. Here, we showcase the use of MALDI iprm-PASEF to provide rapid and accurate annotation coproporphyrin III directly from a bacterial-fungal co-culture between *Glutamicibacter arilaitensis* (strain JB182) and *Penicillium solitum* (strain #12). Additionally, we present a workflow for untargeted iprm-PASEF precursor selection directly in SCiLS Lab, followed by direct export for iprm-PASEF acquisition.

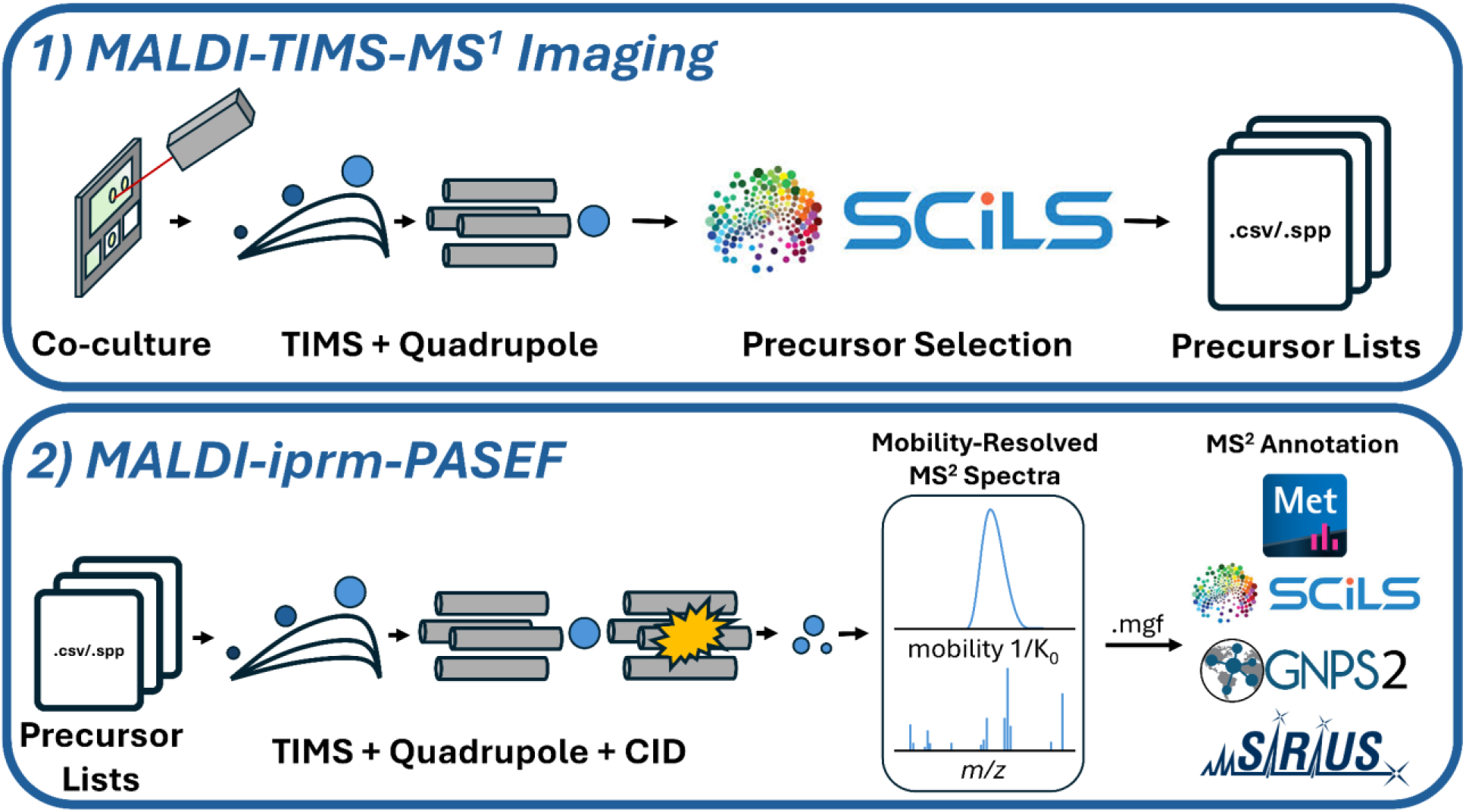

## Introduction

Mass spectrometry imaging (MSI) is a powerful tool for both the targeted and untargeted investigations of the spatial distributions of ions in biological samples.^1^ A notable limitation to MSI is the need for subsequent annotation of the observed MS^1^ level features with complementary datasets such as tandem mass spectrometry (MS^2^) spectra via LC-MS^2^.^2^ The need for these orthogonal datasets often requires a non-trivial amount of additional time and resources to MSI-based projects. Furthermore, the lack of integrated tools to connect these orthogonal datasets and allow for MS^2^ annotation of MSI datasets in an automated fashion is a notable, persistent bottleneck.

Ion mobility spectrometry (IMS) adds dimensionality to MSI datasets, enabling the separation of ions based on their three-dimensional shape and size, and increasing confidence in compound annotation. Trapped ion mobility spectrometry (TIMS) ‘traps’ ions that are propelled by a steady flow of gas against an opposing electric field. Decreasing the strength of the electric field allows the trapped ions to be selectively released from the TIMS cell based on their mobility (K), which is often reported as the inverse reduced mobility (1/K_0_). This value can ultimately be converted to collisional cross section (CCS) for analysis and annotation.^3^ The use of TIMS in MS^2^ experiments, namely parallel accumulation serial fragmentation (PASEF), greatly increases the number of MS^2^ spectra collected within the same timeframe compared to a traditional LC-MS/MS experiment. PASEF technology synchronizes quadrupole isolation with mobility separation allowing for the acquisition of multiple MS^2^ spectra within a single TIMS ramp. This effectively increases MS^2^ scan rates by more than tenfold with minimal impact on sensitivity.^4,5^ PASEF acquisition modes have largely remained exclusive to LC-IMS-MS experiments. However, recent studies and software improvements have implemented parallel reaction monitoring - PASEF (prm-PASEF) in MALDI experiments to enable the collection of multiple MS^2^ spectra within a single MALDI-TIMS scan event.^6,7^ While these studies have showcased the use of MS^2^ data acquired via MALDI imaging prm-PASEF (iprm-PASEF) to perform MS^2^ spectral annotation and molecular networking from MALDI-MSI datasets, they have relied on third party software and prototypic workflows to perform both acquisition and downstream analysis of the data, adding to the complexity of iprm-PASEF data analysis.

MSI has proven to be useful for studying complex microbial systems, allowing researchers to gain information about microbial chemical communication by monitoring the spatial distribution of metabolites across microbial cultures. This can provide valuable information about microbial excretion or retention of metabolites, informing downstream sample preparation and experimental design. Microbial MSI, however, requires the use of complex culture media that often contains salts and other components that can impact ionization in MALDI-MS^1^. Additionally, many microbial metabolites of interest have *m/z* values <1000, and microbial MSI samples tend to be more heterogeneous in surface topology than tissue samples. For these reasons, mitigating matrix/media interference, and making accurate annotations from MALDI-MS^1^ remain a notable challenge. By leveraging the inclusion of TIMS and MS^2^ via MALDI iprm-PASEF, sample interference from matrix/media components can be filtered and accurate annotation can be achieved from a single sample without the need for orthogonal MS experiments. We display the utility of MALDI iprm-PASEF in microbial MSI to accurately annotate the natural product coproporphyrin III, which cannot be accurately identified by MS^2^ analysis alone (Figure S6). We also showcase a vendor-supported workflow for streamlined acquisition and analysis of untargeted MALDI iprm-PASEF datasets using a bacterial-fungal co-culture.

## Experimental Section

### Cultivation of *Glutamicibacter arilaitensis* and *Penicillium solitum*

A frozen stock of *Glutamicibacter arilaitensis* sp. JB182 was removed from -70°C storage. A small amount of the stock was thawed and streaked onto brain heart infusion (BHI) agar (Bacto™, pH 7.4, 1.5% agar) and the plate incubated at RT for 48 hrs to allow colony formation. A single colony was suspended into 5 mL of BHI broth (Bacto™, pH 7.4) and incubated at RT for 24 hr at 200 rpm. The resulting culture was normalized to an optical density (OD_600_) of 0.1. The same day, a pre-normalized (OD_600_ 0.1) frozen stock of *Penicillium solitum* was removed from - 70°C storage and allowed to thaw. In a biosafety hood, 5 µL of the *P. solitum* stock was spotted in the center of a 2mm thick 2.5% cheese curd agar plate (CCA; pH 7.0) and allowed to dry. Next, 5 µL of the normalized *G. arilaitensis* culture was spotted at 1, 2, and 3 cm distances away from the center of the *P. solitum* spot. The plates were incubated for 10 days at RT in the dark until the first and second *G. arilaitensis* colonies (1 cm away) were fully engulfed by *P. solitum*, and the third colony (3 cm away) began coming into contact with the fungal mycelia. All media recipes can be found in the supplementary information (Tables S1 and S2).

### MALDI-MSI sample preparation

Following incubation, the *G. arilaitensis* and *P. solitum* co-cultures along with surrounding agar (∼0.5 cm around the edge of each bacterial colony) were excised from the agar plates using a sterile razor blade and gently transferred to a ground steel MALDI target plate as outlined by Yang et al.^8^ An optical image of the target plate with the colonies was taken prior to matrix application. Standard solutions of coproporphyrins I (1 mM) and III (1 mM) were spotted onto blank squares of CCA agar. A 1:1 mixture of α-cyano-4-hydroxycinnamic acid and 2,5-dihydroxybenzoic acid (1:1 CHCA:DHB) was used as MALDI matrix and applied with a 53 micron stainless steel sieve (Hogentogler & Co, USA). The target plates were allowed to dry at 35°C for 4 hours using a homemade spinning apparatus.^9^ Following drying, excess matrix was removed using compressed air, and 1 µL of saturated red phosphorus in 1:1 ACN/H_2_O was spotted as a calibrant and allowed to dry. Finally, a second optical image of the target plate was taken following matrix application for subsequent MSI acquisition.

### Instrumentation and Software

A full list of software and their respective versions can be found in the supplementary material (Table S3). All data were collected on a timsTOF fleX mass spectrometer (Bruker Daltonics) in positive ion mode from *m/z* 100-1100, and 1/K_0_ 0.4-1.80 Vs/cm^2^. A detailed list of method, laser, and tune parameters for MALDI-TIMS-MS and MALDI-TIMS-MS^2^ acquisition can be found in the supplementary material (Table S4). Our GNPS2 task is public and can be accessed via: https://gnps2.org/status?task=6e2e3794132248aeb2dc6458ba67ea69. All GNPS2, MetaboScape, and SIRIUS query parameters can be found in the supplementary material (Tables S7, S8, and S9, respectively). A prototype tool utilizing the SCiLS Lab Python API to batch export iprm-PASEF MS^2^ data from SCiLS Lab to MASCOT generic format (*.mgf) files can be found at: https://github.com/gtluu/SCiLS_Lab_iprm-PASEF_Exporter.

### Reagents

Coproporphyrin I and III standards were purchased from either Frontier Scientific (Newark, DE), or Santa Cruz Biotechnology, Inc. (Dallas, TX).

## Results and Discussion

### Coproporphyrin Production by *G. arilaitensis*

Previously, we reported that production of the zinc-chelating porphyrin derivative, coproporphyrin III, is upregulated in *G. arilaitensis* when grown in co-culture with *Penicillium solitum.*^10^Coproporphyrin III, however, is one of four naturally occurring constitutional isomers, all of which yield identical MS^2^ fragmentation patterns (Figure S7). As such, confirmation of the correct coproporphyrin isomer produced by *G. arilaitensis* required HPLC retention time analysis using commercial standards and co-injections after acquiring MALDI-MSI, LC-MS ^2^, and NMR data. We postulated that MALDI iprm-PASEF would distinguish the correct coproporphyrin isomer without the need for orthogonal MS and spectroscopic datasets. We sought to test this as a proof-of-principle given that we have already documented the production of zinc-coproporphyrin III by numerous orthogonal methods, including identifying its biosynthetic gene cluster in the producing organism.^10^ Both *G. arilaitensis* and *P. solitum* were grown on cheese curd agar, a nutrient-rich medium that drastically increases spectral complexity during MSI.^11,12^ A prototype MALDI prm-PASEF workflow has been reported to provide enhanced signal-to-noise compared to other TIMS-resolved MS^2^ methods in samples of high complexity, making it advantageous in the context of microbial imaging.^6^ Additionally, to test its capability in an untargeted scenario, we performed MALDI iprm-PASEF on a set of 12 precursors selected from a MALDI-TIMS-MS^1^ dataset acquired from the same sample, prioritized using the statistical tools available within the SCiLS Lab (Bruker Daltonics) software.

### MALDI-TIMS-MS^1^ Data Analysis and iprm-PASEF Precursor Selection

As discussed previously, prm-PASEF is a targeted data-dependent acquisition (DDA) method.^6,7^ We first acquired a survey MALDI-TIMS-MS^1^ image of the prepared *G. arilaitensis* and *P. solitum* co-culture in positive mode, monitoring over 100-1100 *m/z* and 0.40-1.80 V·s/cm^2^ to allow for untargeted analysis in addition to monitoring for coproporphyrin III ([M+H]^+^ 655.2768, calcd. for C_36_H_39_N_4_O_8_ ^+^) (Figure 1A). Upon analysis of the MALDI-TIMS-MS^1^ dataset, *m/z* 655.2726 (1/K_0_ 1.2755 ± 0.02 V·s/cm^2^; mass error = 6.4 ppm) was observed with a spatial distribution consistent with that previously reported for coproporphyrin III production in a *G. arilaitensis* and *P. solitum* co-culture.^10^ To filter matrix/media signals prior to precursor selection, the “Find Discriminating Features (ROC)” tool in SCiLS Lab was performed on the feature list generated with the T-ReX^3^ algorithm using the microbial ROIs as “Class 1” and the matrix/media region as “Class 2” (Table S6). The resulting list yielded 15 features distinct to the *G. arilaitensis, P. solitum*, and/or co-culture regions. From this list, *m/z* values 298.3464, 326.3773, and 672.0338 were present in the matrix/media region with minimal intensity and were manually excluded from the iprm-PASEF precursor list. Another *m/z* at 649.3278 was removed due to it being the corresponding sodiated adduct of another analyte present within our list (*m/z* 627.3479). Three additional features at *m/z* 529.3415, 594.3424, and 608.3601 were manually added to the precursor list due to their spatial association with either *G. arilaitensis* or *P. solitum,* and lack of intensity in the matrix/media region (Table 1). This resulted in the final list of 12 precursors that were ultimately included in the MALDI iprm-PASEF precursor list for MS^2^ acquisition (Figure 1B).

**Table 1.**
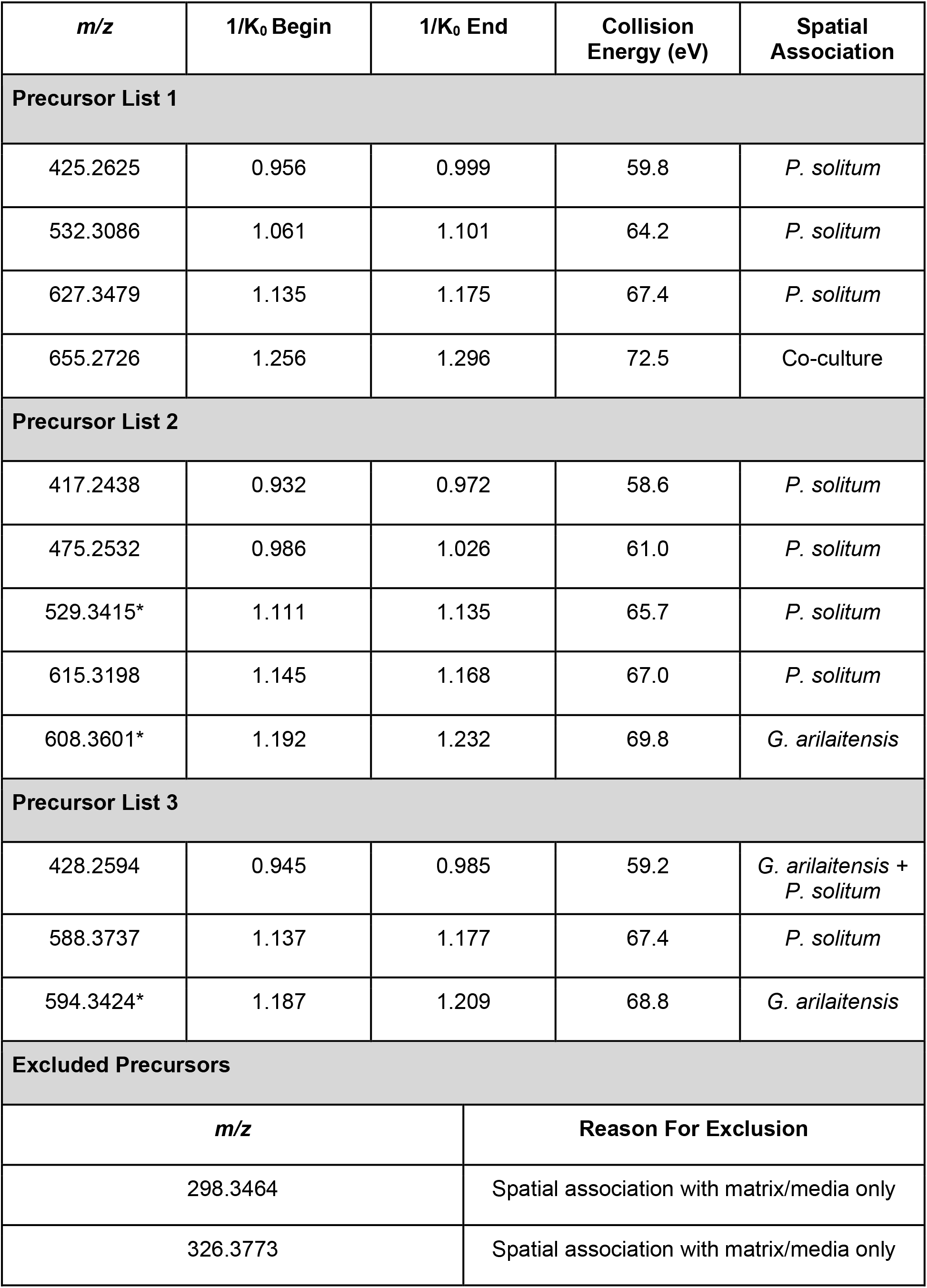
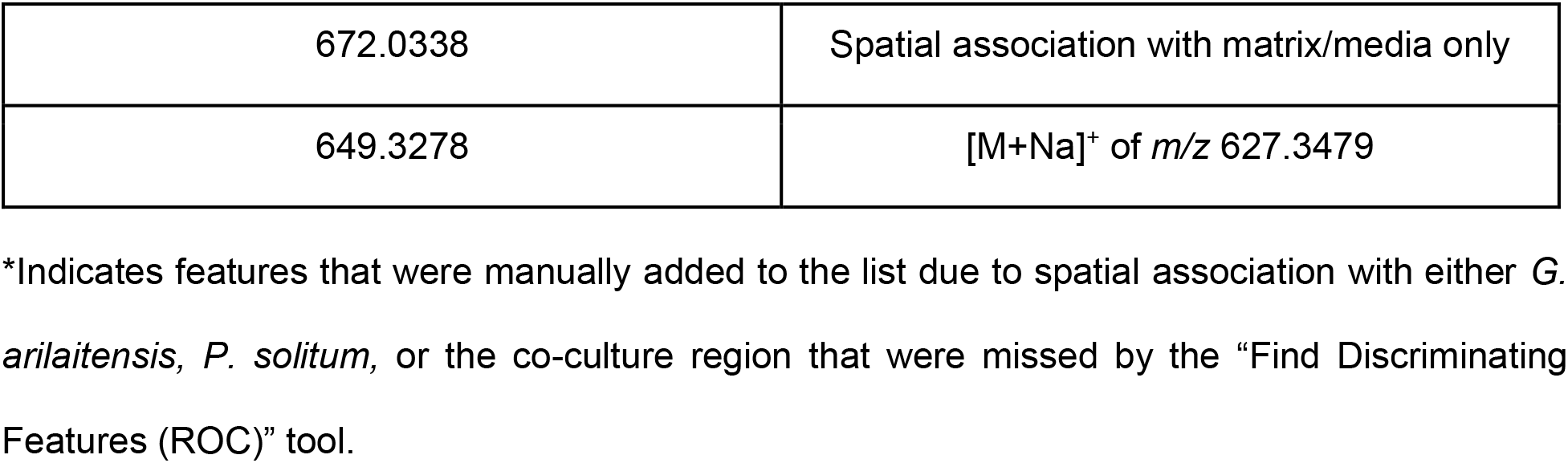
Precursors Selected for MALDI iprm-PASEF Based on ROC Discrimination.

| *m/z* | 1/K<sub>0</sub> Begin | 1/K<sub>0</sub> End | Collision Energy (eV) | Spatial Association |
| --- | --- | --- | --- | --- |
| **Precursor List 1** |  |  |  |  |
| 425.2625 | 0.956 | 0.999 | 59.8 | *P. solitum* |
| 532.3086 | 1.061 | 1.101 | 64.2 | *P. solitum* |
| 627.3479 | 1.135 | 1.175 | 67.4 | *P. solitum* |
| 655.2726 | 1.256 | 1.296 | 72.5 | Co-culture |
| **Precursor List 2** |  |  |  |  |
| 417.2438 | 0.932 | 0.972 | 58.6 | *P. solitum* |
| 475.2532 | 0.986 | 1.026 | 61.0 | *P. solitum* |
| 529.3415* | 1.111 | 1.135 | 65.7 | *P. solitum* |
| 615.3198 | 1.145 | 1.168 | 67.0 | *P. solitum* |
| 608.3601* | 1.192 | 1.232 | 69.8 | *G. arilaitensis* |
| **Precursor List 3** |  |  |  |  |
| 428.2594 | 0.945 | 0.985 | 59.2 | *G. arilaitensis* + *P. solitum* |
| 588.3737 | 1.137 | 1.177 | 67.4 | *P. solitum* |
| 594.3424* | 1.187 | 1.209 | 68.8 | *G. arilaitensis* |
| **Excluded Precursors** |  |  |  |  |
| *m/z* |  | Reason For Exclusion |  |  |
| 298.3464 |  | Spatial association with matrix/media only |  |  |
| 326.3773 |  | Spatial association with matrix/media only |  |  |

|  |  |
| --- | --- |
| 672.0338 | Spatial association with matrix/media only |
| 649.3278 | [M+Na]<sup>+</sup> of *m/z* 627.3479 |

\*Indicates features that were manually added to the list due to spatial association with either *G. arilaitensis*, *P. solitum*, or the co-culture region that were missed by the “Find Discriminating Features (ROC)” tool.

**Figure 1.**
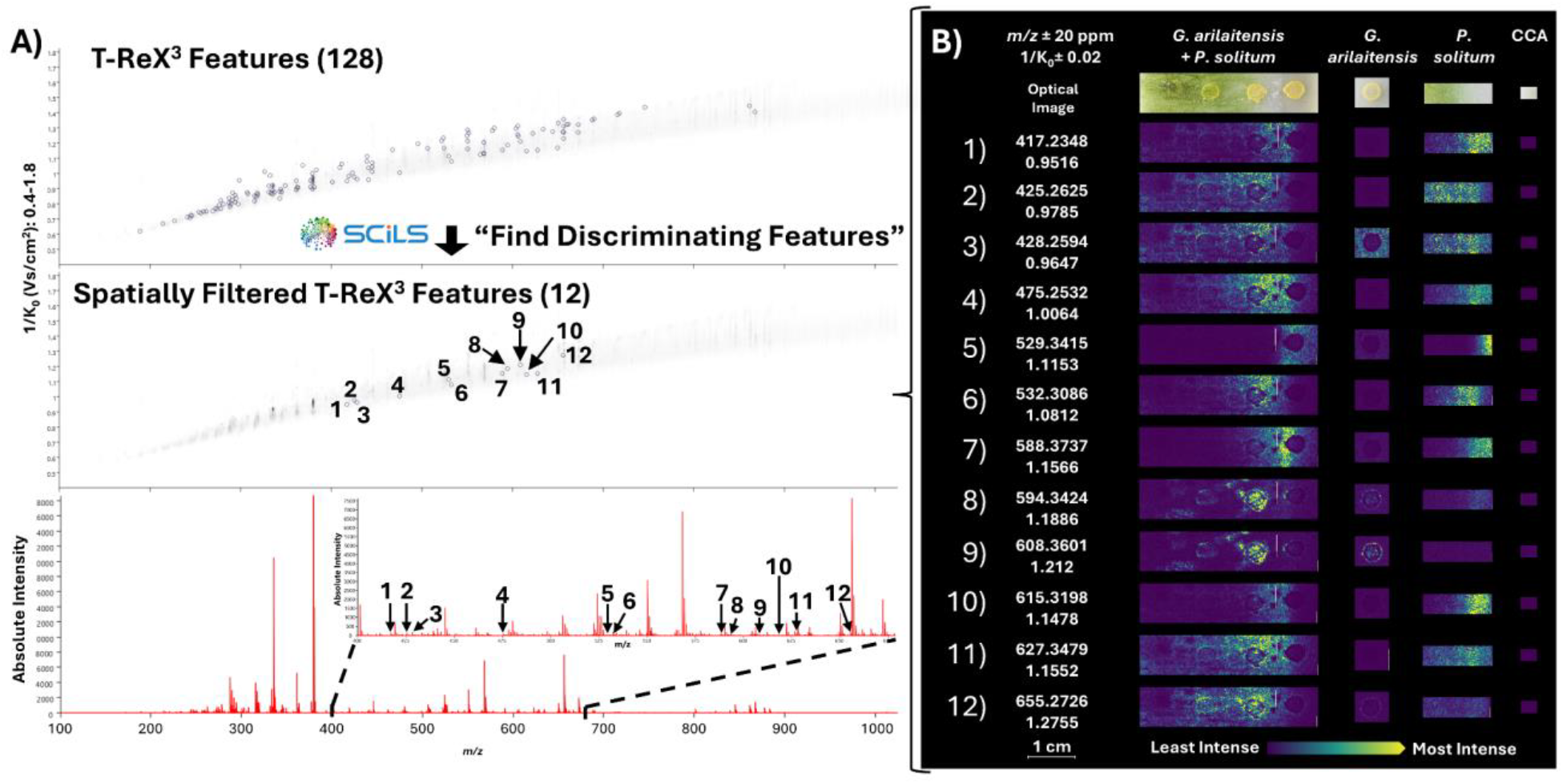
MALDI iprm-PASEF precursor selection from MALDI-TIMS-MS^1^ data in SCiLS Lab. A) The T-ReX^3^ feature finding algorithm identified 128 features using the user-defined parameters (Table S5). The “Find Discriminating Features (ROC)” tool filtered out matrix/media-associated peaks, highlighting features specifically present in the *G. arilaitensis, P. solitum*, and/or co-culture regions. B) Ion images for the twelve MALDI iprm-PASEF precursors selected using SCiLS Lab. All features selected as precursors for MALDI iprm-PASEF are low intensity features relative to matrix/media peaks, highlighting the complexity of microbial MSI.

### iprm-PASEF Data Acquisition

The specific method parameters, experimental setup, and a description of the overall workflow for MALDI iprm-PASEF acquisition are discussed in depth in the supplementary material. MALDI iprm-PASEF allows for the acquisition of multiple MS/MS spectra while allowing for users to manually define collision energy (CE) values for specific precursors or automatically choose CE values based on an interpolation table within a single MALDI-TIMS scan event. Previous analyses have optimized the CE for analytes of interest using either standards or by performing repeated MALDI-TIMS broadband collision induced dissociation (bbCID) experiments to determine optimal CE values for multiple unknown analytes.^6^ MALDI-TIMS-bbCID allows for TIMS-resolved, all-ion fragmentation, but only at a single CE. This approach to optimizing CE in the context of MSI however, requires multiple MALDI-TIMS-bbCID imaging acquisitions, likely exhausting the analyte of interest on a given sample via ablation. This would require access to a separate sample for subsequent MALDI iprm-PASEF acquisition which is not ideal in situations where samples may be difficult to obtain or could experience biological variation. As such, we assessed the performance of MALDI iprm-PASEF using a mobility-interpolated CE table ranging from 35 eV at 0.4 V·s/cm^2^ to 95 eV at 1.80 V·s/cm^2^, enabling the acquisition of a complete MALDI iprm-PASEF workflow from a single sample.

Since prm-PASEF relies on mobility resolution to extract accurate MS^2^ spectra of distinct features, precursors within the same prm-PASEF acquisition cannot have overlapping mobility window ranges. Additionally, mobility windows must have enough time between them to allow for accurate precursor isolation in the quadrupole to prevent MS/MS spectrum “contamination” (i.e. fragments from multiple precursors in the same spectrum). This limits untargeted analyses where there is no guarantee all selected precursors will be mobility-resolved, even if the precursors are mass-resolved. We initially had nine overlapping features within our iprm-PASEF precursor list. To address this, we divided our initial precursor list into three fully mobility-resolved feature lists (Table 1) and multiplexed our MALDI-MSI analysis, splitting each 200×200µm pixel into four 100×100µm quadrants corresponding to the 100 µm laser size (Figure 2A). The first quadrant of each pixel was reserved for the MALDI-TIMS-MS^1^ dataset. The other three quadrants represent the three MALDI iprm-PASEF acquisitions with separate precursor lists. Generating separate mobility-resolved precursor lists from a single feature list can be performed manually in the SCiLS Lab GUI, or automated using the SCiLS lab API, as outlined in the supplemental information.

**Figure 2.**
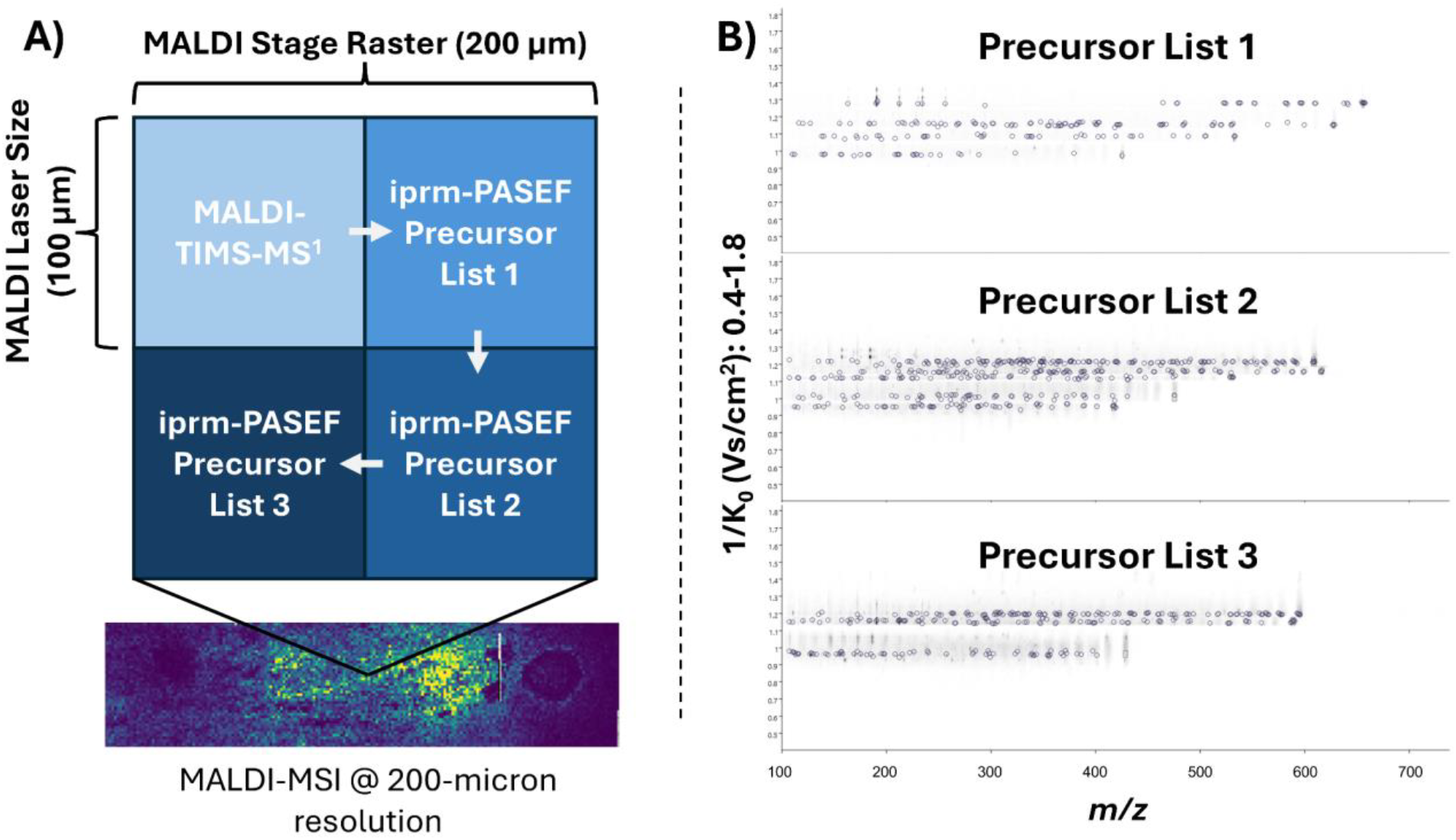
Multiplexed MALDI iprm-PASEF acquisition. A) Schematic representation of a multiplexed MALDI imaging pixel. The first quadrant (no laser offset) is reserved for the MALDI-TIMS-MS^1^ survey scan. The remaining three quadrants (offset by different X,Y increments of 100 µm, or half of the raster width) are utilized for multiple MALDI iprm-PASEF acquisitions of different TIMS-resolved precursor lists. B) Mobility-*m/z* heatmaps of the three acquired MALDI iprm-PASEF precursor lists. SCiLS Lab labels precursor *m/z’s* with dotted line boxes, and TIMS-associated fragment features with circles.

**Figure 3.**
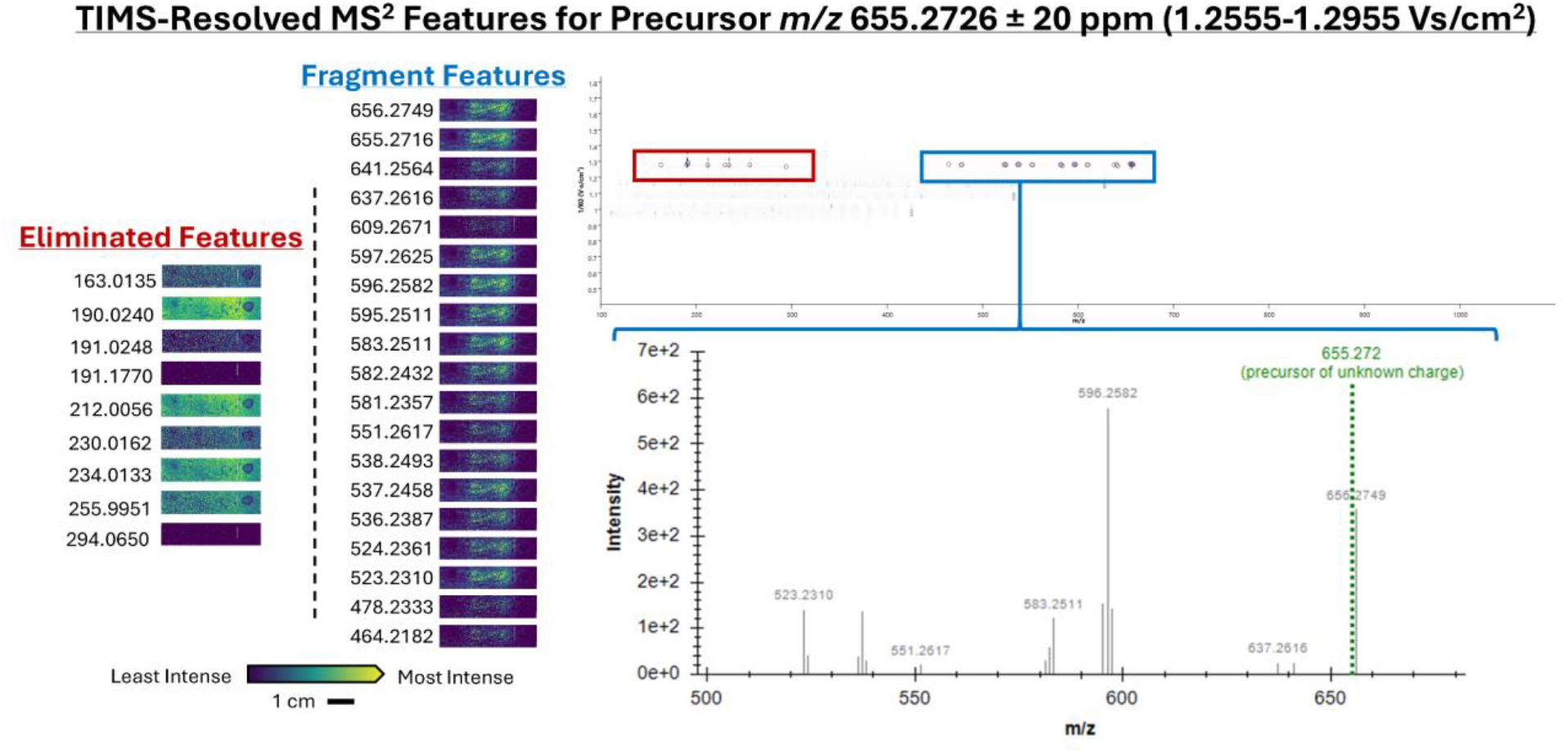
Spatially filtered TIMS-resolved MALDI iprm-PASEF MS^2^ spectrum of *m/z* 655.2726. Fragment features within a defined 1/K_0_ window NOT associated with the precursor of interest can be spatially excluded from the exported MS^2^ spectrum by performing feature co-localization in SCiLS Lab. Note that fragment features associated with the precursor display the same spatial distribution as the precursor of interest.

### Processing and Export of Mobility-Resolved MS^2^ Spectra

Following MALDI iprm-PASEF acquisition, the three flexImaging sequence files (*.mis) were imported separately into SCiLS Lab to yield *.slx files. “Feature Finding” was performed using the T-ReX^3^ algorithm upon import into the software, which automatically labels any *m/z* within each defined 1/K_0_ window as ‘fragments’ belonging to the defined precursor within the same 1/K_0_ window. Nine features selected by the T-ReX^3^ algorithm for *m/z* 655.2726 were not fragments related to the precursor, as determined by examining their spatial distributions. To exclude these unrelated features from the final MS^2^ spectrum, the “Find Values Co-Localized to Feature” workflow was performed, selecting the residual *m/z* 655.2726 precursor as the feature of interest (any feature with a spatial distribution consistent with the defined precursor can be used). This workflow was used to compare fragment ion images to that of *m/z* 655.2726 to ensure that all fragments that did not share the same spatial distribution as the precursor were removed from the final MS^2^ spectrum, which resulted in a spatially-filtered, mobility-resolved MS^2^ spectrum ready for export. Previously, TDF files from MALDI iprm-PASEF analyses had to be uploaded into DataAnalysis (Bruker Daltonics) to generate extracted ion mobilograms (EIMs) for each precursor, followed by manual extraction of averaged MS^2^ spectra. The MS^2^ spectra could subsequently be exported one-by-one as MASCOT generic format (*.mgf) files that required manual addition of “BEGIN IONS” and “END IONS” statements to define the MS^2^ features. We streamlined this process using the SCiLS Lab API to enable direct export of correctly formatted *.mgf files containing either individual MS^2^ spectra, or a consensus .mgf file containing all collected MS^2^ spectra from a single imaging run. The generated *.mgf files can be used directly in downstream software for spectral library searching, including MetaboScape (Bruker Daltonics) and GNPS2.^13^ Lastly, this highlights the importance of considering spatial information for the confirmation of ‘real’ fragments in iprm-PASEF MS^2^ spectra. Workflows utilizing MALDI iprm-PASEF typically export an MS^2^ spectrum that is averaged over a designated mobility range without considering the spatial distribution of the individual features. However, it should be noted that in cases such as low ion abundance, not all precursors and fragments may share completely similar spatial distributions. The presence of unrelated fragments within the same mobility range as the precursor of interest can influence the accuracy of spectral library search results (Figure S2).

### MALDI iprm-PASEF MS^2^ Spectral Annotation

For MS^2^ spectral annotation in GNPS2, separate *.mgf files containing the individual MS^2^ spectra of our 12 analytes of interest were uploaded into the GNPS2 web platform along with a metadata table. The “librarysearch_workflow” was performed with “Top-K” set to 20 to generate a list of the top 20 most likely spectral matches based on cosine similarity between experimental and library MS^2^ spectra. Our GNPS2 library search matched our experimental MS^2^ spectrum for precursor *m/z* 655.2726 with the library MS^2^ spectra of coproporphyrins I and III at cosine similarity scores of 0.96 and 0.89 respectively, indicating a high-confidence annotation of a coproporphyrin-class molecule. Additionally, all 20 GNPS2 library hits for precursor *m/z* 655.2726 were porphyrin-class molecules with cosine similarity scores >0.75. Manual evaluation of the MS^2^ spectrum identified major fragments at *m/z* 637, 596, 537, and 523, consistent with data previously reported in the literature for the coproporphyrins (Figure S1).^10,14^

The MS^2^ spectrum of *m/z* 425.263 yielded several GNPS2 database hits suggestive of a histidine-containing peptide, but no structure assignment was provided for the top GNPS2 library hit (cosine similarity score: 0.86) (Figure S3). MetaboScape Spectral Library annotation identified *m/z* 425.263 as His-Leu-Arg from the “Bruker NIST 2020 MSMS Spectral Library” with relatively low confidence (MS/MS score of 627.80) (Figure S5). Conversely, submitting the spectrum into SIRIUS^15^ gave a calculated annotation for Arg-Leu-His with a SIRIUS score of 94.879% based on fragmentation substructure analysis (Figure S6, Table S10). No other MS^2^ spectra collected in this work yielded GNPS2 library hits with cosine similarity scores >0.77. This could be attributed to low precursor intensity, the use of a generalized CE interpolation, or could indicate a lack of representative database spectra in GNPS2. More in-depth analysis of individual spectra is necessary to confirm this. Lastly, GNPS2 does not currently take CCS into account for classical MS^2^ library searching.

For spectral annotation in MetaboScape, the initial MALDI-TIMS-MS^1^ dataset is imported into the software as a flexImaging sequence file (*.mis) alongside a SCiLS Lab Region Definition (*.srd) file containing coordinate metadata, which can be exported from SCiLS Lab. Feature Finding using the T-ReX^3^ algorithm was performed using the same parameters as the T-ReX^3^ “Feature Finding” in SCiLS Lab to generate a MetaboScape feature table. MetaboScape offers several options for annotation, including from “Target Lists”, which do not contain experimental MS^2^ data by default, and from “Spectral Libraries”, which contain experimental MS^2^ data. Target List annotation combined with SmartFormula annotation can be performed upon import into MetaboScape based on monoisotopic mass, mobility (CCS), adduct *m/z’*s, and chemical formula prediction based on the SmartFormula algorithm. This provides a Level 3 annotation according to the Metabolomics Standard Initiative, and does not guarantee high-confidence compound identification.^16^ For example, when our dataset was initially annotated at the TIMS-MS^1^ level using Natural Product Atlas^17^ as the Target List, *m/z* 655.272 was annotated as thiomarinol D (Calculated [M+H]^+^ 655.2723 for C_31_H_47_N_2_O_9_ S_2_ ^+^), but was later determined to be coproporphyrin (III) based on MS^2^ spectral analysis. This highlighted the importance of MS^2^ spectral matching for confident feature annotation in MSI datasets, even with the implementation of IMS. The individual *.mgf MS^2^ spectra from iprm-PASEF were manually appended to the associated precursor *m/z* in the generated feature table. Spectral Library annotation was performed using the GNPS2 Spectral Library uploaded into MetaboScape as a mass spectrum profile (*.msp) file. Spectral Library query parameters were set to the corresponding thresholds as were set for our GNPS2 query (i.e. an “MS/MS score” of 650 in Metaboscape is analogous to a “Cosine Similarity Score” of 0.65 in GNPS2). The top spectral match in MetaboScape was “Coproporphyrin I or III” from the GNPS2 spectral library (MS/MS score of 869.98). The discrepancy in the top annotations between GNPS2 and MetaboScape can be attributed to the additional query parameters included in the spectral library search function of MetaboScape, and likely differences in their spectral alignment algorithms and criteria. We also manually compared the observed ions to those we have previously reported from fungi grown on CCA and did not find matches other than coproporphyrins.^11,18^

### Confirmation of Coproporphyrin III via TIMS

Since all coproporphyrin isomers yield identical MS/MS spectra, we leveraged the TIMS dimension to accurately annotate the correct isomer. When predicting their CCS values with CCS-Predict Pro in MetaboScape, coproporphyrin I and III were calculated to have identical CCS values at 260.7 Å^2^. Upon repeating the MALDI iprm-PASEF workflow on the *G. arilaitensis* and *P. solitum* co-culture with standards of coproporphyrin I and III, near-baseline resolution was achieved by increasing the TIMS ramp time to 500 ms and narrowing the 1/K_0_ range to 1.29-1.32 Vs/cm^2^ (Figure 4B). A CCS of 267.6 Å^2^ (1/K_0_ = 1.31) was observed for the coproporphyrin III standard and *m/z* 655.27 in the co-culture MSI region, resulting in 2.6% error with respect to the value calculated by CCS-Predict Pro. The combination of ion mobility and MS^2^ spectral annotation allowed for rapid, high confidence annotation of coproporphyrin III from a fungal-bacterial co-culture without further cultivation, extraction, or chromatographic studies (Figure 4).

**Figure 4.**
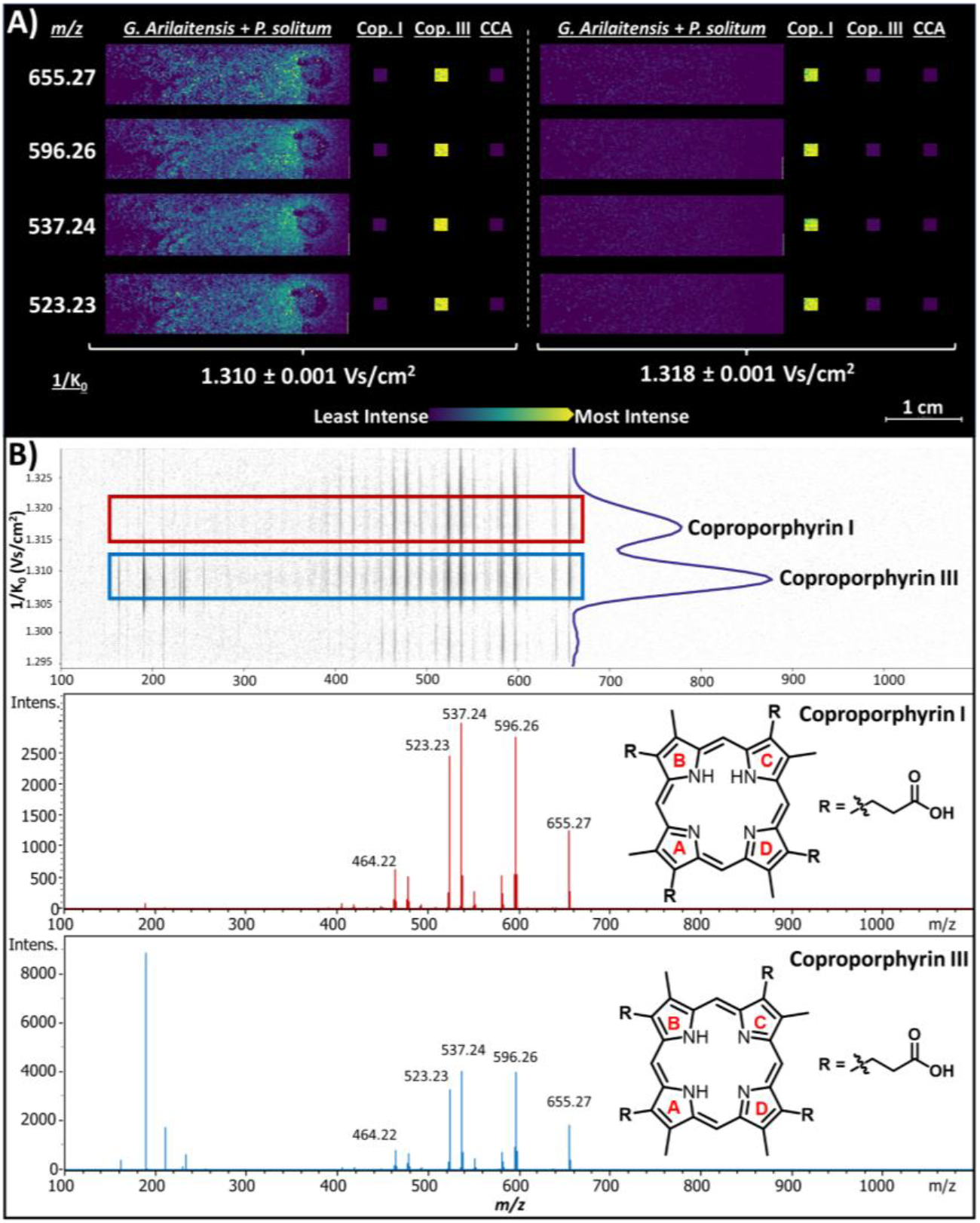
Confirmation of coproporphyrin III in co-culture using MALDI iprm-PASEF. A) Ion images for major fragments of coproporphyrin, resolved only by 1/K_0_. Major fragments for *m/z* 655.27 in co-culture match the mobility range associated with the coproporphyrin III standard, and are consistent with previously reported spatial distributions of coproporphyrin III production by *G. arilaitensis* in co-culture. B) Mobility-resolved MS^2^ spectra and structures of coproporphyrin I (red) and III (blue). Key abbreviations: “cop” = coproporphyrin; “CCA” = cheese curd agar.

## Conclusions

This work highlights a vendor-supported workflow for MALDI iprm-PASEF utilizing commercially available software for both acquisition and analysis. Furthermore, we explored the current capabilities of MALDI iprm-PASEF for untargeted molecular annotation in a fungal-bacterial co-culture grown on solid agar. We acquired MS^2^ spectra for 12 separate precursors on a single co-culture sample and highlighted the utility of multiplexed MSI analysis to maximize MS^2^ spectral collection in a case where not all precursors of interest were fully TIMS-resolved. Excitingly, we demonstrated that combining TIMS and MS^2^ data could distinguish between coproporphyrin analogs, highlighting the capability of MALDI iprm-PASEF to accurately and rapidly facilitate natural product annotation directly from single cultures. Our previous work in annotating coproporphyrin III from *G. arilaitensis* in co-culture required an additional 7-10 days for large-scale culturing and extraction, followed by orthogonal validation using HPLC, LC-MS^2^, and NMR. The analysis presented here required no additional culturing, and confirmation of coproporphyrin III via MALDI iprm-PASEF was performed within ∼3 hours, highlighting the considerable time and resources saved by employing iprm-PASEF for MALDI-MSI annotation. Lastly, we provide an open-source script utilizing the SCiLS Lab API to streamline the generation and export of multiple .mgf files from MALDI iprm-PASEF datasets analyzed in SCiLS Lab, enabling their use in downstream annotation software such as GNPS2. We anticipate this workflow will benefit the broader microbial MSI and NP communities, and is a stepping stone for untargeted spatial metabolomics of complex microbial systems.

## Acknowledgements

This work was supported by the National Institute of General Medical Sciences of the NIH award R21GM148870 (LMS) and by National Science Foundation (NSF) grants IOS-2220510 (LMS) and NSF Graduate Research Fellowship Program (RAS).

## Data Availability

All MALDI iprm-PASEF data in this study are available under CC0 1.0 Universal License as raw (*.tdf) and open source (*.imzML) data formats. MassIVE accession: MSV000097837; https://doi.org/doi:10.25345/C5JH3DF50.

## Supporting Information

The Supporting Information is available free of charge at https://pubs.acs.org/doi/XXX.

- Media recipes, and all instrument, method, and database query parameters used in the collection and analysis of the data in this manuscript, additional MALDI iprm-PASEF data and database query results that support the identification of coproporphyrin III, and the annotation of a small molecule peptide, and a detailed description of experimental design, data collection, and data analysis of MALDI iprm-PASEF data.

## References

(1) Spraker, J. E.; Luu, G. T.; Sanchez, L. M. Imaging Mass Spectrometry for Natural Products Discovery: A Review of Ionization Methods. Nat. Prod. Rep. 2019. 10.1039/c9np00038k.

(2) McAtamney, A.; Heaney, C.; Lizama-Chamu, I.; Sanchez, L. M. Reducing Mass Confusion over the Microbiome. Anal. Chem. 2023. 10.1021/acs.analchem.3c02408.

(3) Gabelica, V.; Shvartsburg, A. A.; Afonso, C.; Barran, P.; Benesch, J. L. P.; Bleiholder, C.; Bowers, M. T.; Bilbao, A.; Bush, M. F.; Campbell, J. L.; Campuzano, I. D. G.; Causon, T.; Clowers, B. H.; Creaser, C. S.; De Pauw, E.; Far, J.; Fernandez-Lima, F.; Fjeldsted, J. C.; Giles, K.; Groessl, M.; Hogan, C. J., Jr; Hann, S.; Kim, H. I.; Kurulugama, R. T.; May, J. C.; McLean, J. A.; Pagel, K.; Richardson, K.; Ridgeway, M. E.; Rosu, F.; Sobott, F.; Thalassinos, K.; Valentine, S. J.; Wyttenbach, T. Recommendations for Reporting Ion Mobility Mass Spectrometry Measurements. Mass Spectrom. Rev. 2019, 38 (3), 291–320.

(4) Vasilopoulou, C. G.; Sulek, K.; Brunner, A.-D.; Meitei, N. S.; Schweiger-Hufnagel, U.; Meyer, S. W.; Barsch, A.; Mann, M.; Meier, F. Trapped Ion Mobility Spectrometry and PASEF Enable in-Depth Lipidomics from Minimal Sample Amounts. Nat. Commun. 2020, 11 (1), 331.

(5) Meier, F.; Beck, S.; Grassl, N.; Lubeck, M.; Park, M. A.; Raether, O.; Mann, M. Parallel Accumulation-Serial Fragmentation (PASEF): Multiplying Sequencing Speed and Sensitivity by Synchronized Scans in a Trapped Ion Mobility Device. J. Proteome Res. 2015, 14 (12), 5378–5387.

(6) Wolf, C.; Behrens, A.; Brungs, C.; Mende, E. D.; Lenz, M.; Piechutta, P. C.; Roblick, C.; Karst, U. Mobility-Resolved Broadband Dissociation and Parallel Reaction Monitoring for Laser Desorption/ionization-Mass Spectrometry - Tattoo Pigment Identification Supported by Trapped Ion Mobility Spectrometry. Anal. Chim. Acta 2023, 1242 (340796), 340796.

(7) Heuckeroth, S.; Behrens, A.; Wolf, C.; Fütterer, A.; Nordhorn, I. D.; Kronenberg, K.; Brungs, C.; Korf, A.; Richter, H.; Jeibmann, A.; Karst, U.; Schmid, R. On-Tissue Dataset-Dependent MALDI-TIMS-MS2 Bioimaging. Nat. Commun. 2023, 14 (1), 7495.

(8) Yang, J. Y.; Phelan, V. V.; Simkovsky, R.; Watrous, J. D.; Trial, R. M.; Fleming, T. C.; Wenter, R.; Moore, B. S.; Golden, S. S.; Pogliano, K.; Dorrestein, P. C. Primer on Agar-Based Microbial Imaging Mass Spectrometry. J. Bacteriol. 2012, 194 (22), 6023–6028.

(9) Lusk, H. J.; Levy, S. E.; Bergsten, T. M.; Burdette, J. E.; Sanchez, L. M. Home-Built Spinning Apparatus for Drying Agarose-Based Imaging Mass Spectrometry Samples. J. Am. Soc. Mass Spectrom. 2022, 33 (7), 1325–1328.

(10) Cleary, J. L.; Kolachina, S.; Wolfe, B. E.; Sanchez, L. M. Coproporphyrin III Produced by the Bacterium Glutamicibacter Arilaitensis Binds Zinc and Is Upregulated by Fungi in Cheese Rinds. mSystems 2018, 3 (4). 10.1128/msystems.00036-18.

(11) Luu, G. T.; Little, J. C.; Pierce, E. C.; Morin, M.; Ertekin, C. A.; Wolfe, B. E.; Baars, O.; Dutton, R. J.; Sanchez, L. M. Metabolomics of Bacterial-Fungal Pairwise Interactions Reveal Conserved Molecular Mechanisms. Analyst 2023. 10.1039/d3an00408b.

(12) Cleary, J. L.; Luu, G. T.; Pierce, E. C.; Dutton, R. J.; Sanchez, L. M. BLANKA: An Algorithm for Blank Subtraction in Mass Spectrometry of Complex Biological Samples. J. Am. Soc. Mass Spectrom. 2019, 30 (8), 1426–1434.

(13) Wang, M.; Carver, J. J.; Phelan, V. V.; Sanchez, L. M.; Garg, N.; Peng, Y.; Nguyen, D. D.; Watrous, J.; Kapono, C. A.; Luzzatto-Knaan, T.; Porto, C.; Bouslimani, A.; Melnik, A. V.; Meehan, M. J.; Liu, W.-T.; Crüsemann, M.; Boudreau, P. D.; Esquenazi, E.; Sandoval-Calderón, M.; Kersten, R. D.; Pace, L. A.; Quinn, R. A.; Duncan, K. R.; Hsu, C.-C.; Floros, D. J.; Gavilan, R. G.; Kleigrewe, K.; Northen, T.; Dutton, R. J.; Parrot, D.; Carlson, E. E.; Aigle, B.; Michelsen, C. F.; Jelsbak, L.; Sohlenkamp, C.; Pevzner, P.; Edlund, A.; McLean, J.; Piel, J.; Murphy, B. T.; Gerwick, L.; Liaw, C.-C.; Yang, Y.-L.; Humpf, H.-U.; Maansson, M.; Keyzers, R. A.; Sims, A. C.; Johnson, A. R.; Sidebottom, A. M.; Sedio, B. E.; Klitgaard, A.; Larson, C. B.; Boya P, C. A.; Torres-Mendoza, D.; Gonzalez, D. J.; Silva, D. B.; Marques, L. M.; Demarque, D. P.; Pociute, E.; O’Neill, E. C.; Briand, E.; Helfrich, E. J. N.; Granatosky, E. A.; Glukhov, E.; Ryffel, F.; Houson, H.; Mohimani, H.; Kharbush, J. J.; Zeng, Y.; Vorholt, J. A.; Kurita, K. L.; Charusanti, P.; McPhail, K. L.; Nielsen, K. F.; Vuong, L.; Elfeki, M.; Traxler, M. F.; Engene, N.; Koyama, N.; Vining, O. B.; Baric, R.; Silva, R. R.; Mascuch, S. J.; Tomasi, S.; Jenkins, S.; Macherla, V.; Hoffman, T.; Agarwal, V.; Williams, P. G.; Dai, J.; Neupane, R.; Gurr, J.; Rodríguez, A. M. C.; Lamsa, A.; Zhang, C.; Dorrestein, K.; Duggan, B. M.; Almaliti, J.; Allard, P.-M.; Phapale, P.; Nothias, L.-F.; Alexandrov, T.; Litaudon, M.; Wolfender, J.-L.; Kyle, J. E.; Metz, T. O.; Peryea, T.; Nguyen, D.-T.; VanLeer, D.; Shinn, P.; Jadhav, A.; Müller, R.; Waters, K. M.; Shi, W.; Liu, X.; Zhang, L.; Knight, R.; Jensen, P. R.; Palsson, B.Ø.; Pogliano, K.; Linington, R. G.; Gutiérrez, M.; Lopes, N. P.; Gerwick, W. H.; Moore, B. S.; Dorrestein, P. C.; Bandeira, N. Sharing and Community Curation of Mass Spectrometry Data with Global Natural Products Social Molecular Networking. Nat. Biotechnol. 2016, 34 (8), 828–837.

(14) Perez-Ortiz, G.; Sidda, J. D.; Peate, J.; Ciccarelli, D.; Ding, Y.; Barry, S. M. Production of Copropophyrin III, Biliverdin and Bilirubin by the Rufomycin Producer, Streptomyces Atratus. Front. Microbiol. 2023, 14, 1092166.

(15) Dührkop, K.; Fleischauer, M.; Ludwig, M.; Aksenov, A. A.; Melnik, A. V.; Meusel, M.; Dorrestein, P. C.; Rousu, J.; Böcker, S. SIRIUS 4: A Rapid Tool for Turning Tandem Mass Spectra into Metabolite Structure Information. Nat. Methods 2019, 16 (4), 299–302.

(16) Sumner, L. W.; Amberg, A.; Barrett, D.; Beale, M. H.; Beger, R.; Daykin, C. A.; Fan, T. W.-M.; Fiehn, O.; Goodacre, R.; Griffin, J. L.; Hankemeier, T.; Hardy, N.; Harnly, J.; Higashi, R.; Kopka, J.; Lane, A. N.; Lindon, J. C.; Marriott, P.; Nicholls, A. W.; Reily, M. D.; Thaden, J. J.; Viant, M. R. Proposed Minimum Reporting Standards for Chemical Analysis Chemical Analysis Working Group (CAWG) Metabolomics Standards Initiative (MSI): Chemical Analysis Working Group (CAWG) Metabolomics Standards Initiative (MSI). Metabolomics 2007, 3 (3), 211–221.

(17) van Santen, J. A.; Jacob, G.; Singh, A. L.; Aniebok, V.; Balunas, M. J.; Bunsko, D.; Neto, F. C.; Castaño-Espriu, L.; Chang, C.; Clark, T. N.; Cleary Little, J. L.; Delgadillo, D. A.; Dorrestein, P. C.; Duncan, K. R.; Egan, J. M.; Galey, M. M.; Haeckl, F. P. J.; Hua, A.; Hughes, A. H.; Iskakova, D.; Khadilkar, A.; Lee, J.-H.; Lee, S.; LeGrow, N.; Liu, D. Y.; Macho, J. M.; McCaughey, C. S.; Medema, M. H.; Neupane, R. P.; O’Donnell, T. J.; Paula, J. S.; Sanchez, L. M.; Shaikh, A. F.; Soldatou, S.; Terlouw, B. R.; Tran, T. A.; Valentine, M.; van der Hooft, J. J. J.; Vo, D. A.; Wang, M.; Wilson, D.; Zink, K. E.; Linington, R. G. The Natural Products Atlas: An Open Access Knowledge Base for Microbial Natural Products Discovery. ACS Cent Sci 2019, 5 (11), 1824–1833.

(18) Pierce, E. C.; Morin, M.; Little, J. C.; Liu, R. B.; Tannous, J.; Keller, N. P.; Pogliano, K.; Wolfe, B. E.; Sanchez, L. M.; Dutton, R. J. Bacterial-Fungal Interactions Revealed by Genome-Wide Analysis of Bacterial Mutant Fitness. Nat Microbiol 2021, 6 (1), 87–102.

(19) Ross, D. H.; Cho, J. H.; Xu, L. Breaking down Structural Diversity for Comprehensive Prediction of Ion-Neutral Collision Cross Sections. Anal. Chem. 2020, 92 (6), 4548–4557.

