## Supplementary Information for "MALDI-TIMS-MS^2^ Imaging and Annotation of Natural Products in Fungal-Bacterial Co-Culture"

**Table S1.** Bacto™ Brain Heart Infusion (BHI) agar recipe………………………………................... S1
**Table S2.** Cheese Curd agar (CCA) recipe…………………………........................................... S2
**Table S3.** Software information……………………………………………………………………… S3
**Table S4.** Instrument parameters for MALDI-TIMS-MSI…………………………………………….. S4-S5
**Workflow S1.** MALDI iprm-PASEF……………………………………………………………………. S6-S13
**Table S5.** T-ReX$^3$ feature finding algorithm parameters……………………..……………………… S7
**Table S6.** “Find Discriminating Features (ROC)” parameters……………………………………….. S8
**Table S7.** “Find Values Co-Localized to Feature” parameters……..………………………… S11
**Table S8.** GNPS2 library search parameters…………………………..………………………….. S14
**Figure S1.** Confirmation of coproporphyrin III in co-culture by MS$^2$ and TIMS resolution………... S15
**Figure S2.** GNPS2 annotation of non-spatially filtered MS$^2$ for *m/z* 655.27……………………….. S16
**Figure S3.** Top GNPS2 library annotations for *m/z* 425.36………………………………………… S17
**Table S9.** Metaboscape spectral library annotation method parameters……………..…………... S18
**Figure S4.** Metaboscape spectral library search results for *m/z* 655.273………………………… S19
**Figure S5.** Metaboscape spectral library search results for *m/z* 425.262………………………… S20
**Table S10.** SIRIUS Compute parameters……………………………………………………………... S21
**Figure S6.** SIRIUS search results and MS/MS fragmentation tree……………………………… S22
**Figure S7.** EIM and averaged MS/MS spectra of coproporphyrin III standards…...………………. S23
**Figure S8.** MALDI-TIMS-MS1 ion images of previously reported isotopologue *m/z*’s for zinc-coproporphyrin……………………………………………………………………………………… S24

**Table S1. Bacto™ Brain Heart Infusion (BHI) Agar Recipe (per 1L of MilliQ Water)**

| Component | Amount (g) |
|---|---|
| Bacto™ Brain Heart Infusion | 37 |
| Agar | 15 |
| pH 7.4 (adjust with 1M NaOH) | |

*Autoclaved at 121°C for 15 minutes per manufacturer's instructions.

**Table S2. Cheese Curd Agar (CCA) Recipe (per 1L of MilliQ Water)**

| Component | Amount (g) |
|---|---|
| Lyophilized cheese curds (Jasper Hills) | 25 |
| Xanthan gum | 5 |
| NaCl | 30 |
| Agar | 17 |
| pH 7.0 (adjust with 1M NaOH) | |

*Autoclaved dry at 121°C for 15 minutes with no dry time. Longer cycle times cause the cheese medium to curdle.

**Table S3. Software Information**

| Software | Version |
|---|---|
| timsControl | 6.0.3_abe6ecc8_1 |
| flexImaging | 7.5 |
| SCiLS Lab | 2025b Pro |
| DataAnalysis | 6.1 (Build 213.27.0)(64-bit) |
| MetaboScape | 2024b |
| GNPS2 | Release 2025.02.12 |
| SIRIUS | 5.8.6 |

**Table S4. Instrument Parameters for MALDI-TIMS-MSI**

| **MS Settings** | | **MS[1]** | **MS[2] (UT)*** | **MS[2] (T)**** |
|---|---|---|---|---|
| Scan range | | 100-1100 *m/z* | 100-1100 *m/z* | 100-1100 *m/z* |
| Ion polarity | | positive | positive | positive |
| Scan mode | | MS | prm-PASEF | prm-PASEF |
| **Tune** | | | | |
| Transfer | MALDI Plate Offset | 50.0 V | 50.0 V | 50.0 V |
| | Deflection 1 Delta | 70.0 V | 70.0 V | 70.0 V |
| | Funnel 1 RF | 225.0 Vpp | 225.0 Vpp | 225.0 Vpp |
| | isCID Energy | 0.0 eV | 0.0 eV | 0.0 eV |
| | Funnel 2 RF | 250.0 Vpp | 250.0 Vpp | 250.0 Vpp |
| | Multipole RF | 350.0 Vpp | 350.0 Vpp | 350.0 Vpp |
| Collision Cell | Collision Energy | 10.0 eV | 10.0 eV | 10.0 eV |
| | Collision RF | 700.0 Vpp | 700.0 Vpp | 700.0 Vpp |
| Quadrupole | Ion Energy | 5.0 eV | 5.0 eV | 5.0 eV |
| | Low Mass | 100.00 *m/z* | 100.00 *m/z* | 100.00 *m/z* |
| Focus Pre TOF | Transfer Time | 65.0 µs | 65.0 µs | 65.0 µs |
| | Pre Pulse Storage | 4.0 µs | 4.0 µs | 4.0 µs |
| **Laser** | | | | |
| Application | | Imaging 100µm | Imaging 100µm | Imaging 100µm |
| Power Boost | | 3.0% | 3.0% | 3.0% |
| Scan Range | | X= 26.0 µm, Y= 26.0 µm | X= 26.0 µm, Y= 26.0 µm | X= 26.0 µm, Y= 26.0 µm |

| | | | |
|---|---|---|---|
| Resulting Field Size | X= 100.0 µm, Y= 100.0 µm | X= 100.0 µm, Y= 100.0 µm | X= 100.0 µm, Y= 100.0 µm |
| Laser Power | 50% | 50% | 50% |
| Bursts | 1 | 1 | 1 |
| Shots | 1000 | 1000 | 1000 |
| Frequency | 5000 Hz | 5000 Hz | 5000 Hz |
| **TIMS** | | | |
| $1/K_0$ start | 0.40 | 0.40 | 1.29 |
| $1/K_0$ end | 1.80 | 1.80 | 1.32 |
| Ramp time | 200.0 ms | 200.0 ms | 500 |
| Accumulation time | 201.8 ms | 201.8 ms | 201.8 |
| Duty cycle | 100.0% | 100.0% | 40.36% |
| Ramp Rate | 4.81 Hz | 4.81 Hz | 1.93 Hz |
| Δt1 (deflection transfer ->capillary exit) | -20.0 V | -20.0 V | -20.0 V |
| Δt2 (deflection transfer ->deflection discard) | -120.0 V | -120.0 V | -120.0 V |
| Δt3 (funnel 1 in ->deflection transfer) | 70 V | 70 V | 70 V |
| Δt4 (accumulation trap ->funnel 1 in) | 100 V | 100 V | 100 V |
| Δt5 (accumulation exit->accumulation transfer) | 0.0 V | 0.0 V | 0.0 V |
| Δt6 (ramp start->accumulation exit) | 100.0 V | 100.0 V | 100.0 V |
| Collision cell in | 220.0 V | 220.0 V | 220.0 V |

| | | | |
|---|---|---|---|
| Lock accumulation to mobility range | Yes | Yes | Yes |

*UT = untargeted

**T = targeted for coproporphyrins

**<u>Workflow S1. MALDI iprm-PASEF</u>**

1. TIMS-$MS^1$ image acquisition
2. Feature finding and construction of feature table
3. Precursor selection in SCiLS Lab
4. iprm-PASEF data acquisition
5. Generation and export of $MS^2$ spectra
6. $MS^2$ Spectral annotation

#### 1) TIMS-$MS^1$ image acquisition

MALDI-TIMS-MS positive ion mode data were collected on a timsTOF fleX (Bruker Daltonics GmbH & Co. KG) using timsControl (Version 6.0.3_abe6ecc8_1) and flexImaging (Version 7.5). The mass range was set to *m/z* 100-1100 in the 'MS' scan mode with 'TIMS' turned on. For our untargeted workflow, the $1/K_0$ range was set to 0.40-1.80 V·s/$cm^2$ with a ramp time of 200.0 ms, an accumulation time of 201.8 ms (set by the duty cycle), and the duty cycle locked at 100%. The M5 laser was selected for analysis, using the default "Imaging 100µm" application setting. The raster size was set to 200 µm in flexImaging and laser size was set to 100 µm in timsControl to allow for multiplexed data acquisition (i.e. collection of $MS^1$ and $MS^2$ spectra within the same MALDI MSI pixel). Prior to acquisition, calibration for *m/z* was performed using phosphorus red in 'Enhanced Quadratic' mode with TIMS turned 'ON'. Mobility calibration was performed using ESI (MALDI 'OFF') with Agilent Tune Mix in 'Linear' mode. The imaging data (.mis file) were loaded into SCiLS Lab (Version 2025b Pro) and converted to a *.slx file upon import into the software.

#### 2) 'Feature Finding' and Construction of Feature Table(s)

The 'Feature Finding' tool is used to automatically construct a processed feature list and can be used upon dataset import into SCiLS Lab, or performed post-import by navigating to the "Tools" tab and selecting "Feature finding". For this study feature-finding was performed upon import into SCiLS Lab using root mean square (RMS) intensity normalization followed by the T-ReX[3] algorithm (parameters below). This generates a global feature list, which often contains matrix or media peaks at high intensities. This interferes with automated, intensity-based precursor selection via use of the SCiLS Lab API. As such, parsing out signals of interest using the statistical tools in SCiLS Lab may be appropriate. The ‘Find discriminating features (ROC)’ tool was used to make a feature list consisting only of features that were associated with the *G. arilaitensis, P. solitum,* and/or co-culture regions of the dataset and not with the cheese curd agar blank. This ensured that the limited collection of $MS^2$ data via iprm-PASEF wasn’t wasted on non-microbial features. Specific parameters for Receiver Operating Characteristic (ROC) analysis can be found in Table S6 below. A detailed explanation of ROC analysis can be found in the SCiLS™ Lab Version 2025b / Release 13.01 user manual (Section 5.6, p. 103-104).

**Table S5. T-Rex$^3$ Feature Finding Algorithm Parameters**

| **T-Rex$^3$ parameters** | |
|---|---|
| **Algorithm** | T-Rex$^3$ (TIMS) |
| **Spatial averaging** | 2x2 |
| **Coverage** | 100% |
| **Maximum features** | 2000 |
| **Polarity** | Positive |
| **Intensity threshold** | Relative - 0.30% |
| **m/z interval width** | Custom - ± 20.000 ppm |
| **Mobility width** | 1/K0 - ± 0.020 V·s / cm$^2$ |
| **CCS extraction** | Extract CCS images automatically |
| **Feature list name** | T-ReX$^3$ features |

**Table S6. “Find Discriminating Features (ROC)” parameters**

| | |
|---|---|
| **Data** | |
| **Normalization** | Root Mean Square |
| **Class 1** | • “Co_1 - Co_4” = *G. arilaitensis* + *P. solitum* co-culture<br>• “Gluta” = *G. arilaitensis*<br>• “P_solitum” = *P. solitum* |
| **Class 2** | “CCA” = 2.5% cheese curd agar |
| • Work on individual spectra | |
| **Feature List** | T-ReX$^3$ Features |
| **Parameters** | |
| **Spectra to Work on** | Random subset |
| **Subset size*** | 100 spectra |

*Receiver Operating Characteristic (ROC) relies on class 1 and class 2 containing a similar number of spectra for accurate results. Since class 1 is made up of six regions and class 2 consists of only one, a random subset of 100 spectra was used to perform the ROC discrimination.

### 3) Precursor Selection In SCiLS Lab

Selection of precursors uses the “Export iprm-PASEF parameters” option in the Feature Table window of SCiLS Lab. The generated table showcases all measured *m/z* values and their corresponding 1/$K_0$ ranges for the selected feature list. Precursors for iprm-PASEF can be selected manually using the “Export iprm-PASEF parameters” module. If multiple features are selected that have overlapping 1/$K_0$ ranges, the “1/$K_0$ start” and “1/$K_0$ end” values will be highlighted in red within the table. iprm-PASEF depends on full resolution of ions via TIMS for successful collection of parallel $MS^2$ spectra within a single scan. Therefore, it is imperative that the features selected for fragmentation within a single iprm-PASEF acquisition do not have overlapping 1/$K_0$ ranges. Features of interest with overlapping mobility ranges must be fragmented in separate iprm-PASEF acquisitions on the same region of interest (ROI). This can be achieved by appropriately applying an offset to the sample carrier stage in flexImaging and adjusting the MALDI laser size in timsControl. The list of precursor *m/z* values and mobility ranges can then be exported as either a SCiLS iprm-PASEF parameter file (*.spp) or as a *.csv file that can be subsequently imported into timsControl for iprm-PASEF acquisition. We used the SCiLS Lab API in Python to automatically sort and prioritize features based on intensity, perform deisotoping to prevent selection of isotopologues, filter them based on mobility range overlap, and ultimately generated three iprm-PASEF *.csv precursor lists with non-overlapping features. Features were prioritized based on their mean intensity across all pixels/spectra.

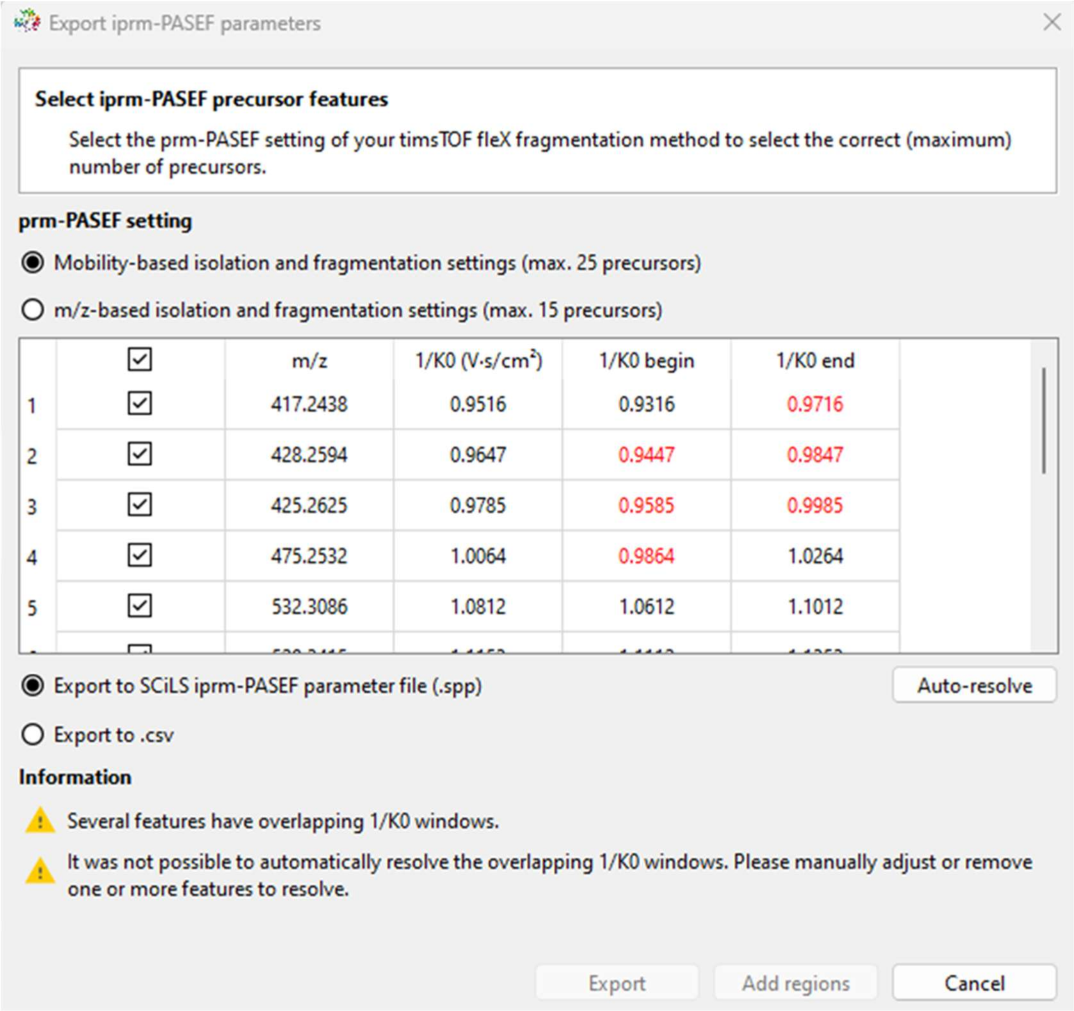

#### 4) iprm-PASEF data acquisition

MALDI iprm-PASEF (MALDI-tims-$MS^2$) data were collected in positive ion mode with the scan mode set to "prm-PASEF". All applicable method parameters and instrument settings remained identical to the aforementioned MALDI-TIMS-$MS^1$ method. Under the "MS/MS" tab in the timsControl method, the generated iprm-PASEF parameter files (.csv) were loaded into the "prm-PASEF windows" module. It should be noted that the allowed $1/K_0$ window is inversely correlated to the ramp time; i.e. higher ramp times allow for narrower $1/K_0$ windows. Therefore, the assigned $1/K_0$ window should reflect the resolution allotted by the ramp time. timsControl will give an error when the assigned $1/K_0$ window for prm-PASEF is too narrow for the given method parameters. Collision energies for $MS^2$ fragmentation were interpolated based on mobility isolation, ranging from 35 eV at 0.4 $V{\cdot}s/cm^2$ to 95 eV at 1.80 $V{\cdot}s/cm^2$. The "Isolation Width Settings" were set to the default values, interpolating from ± 2.00 Da at 700 *m/z* to ± 3.00 Da at 800 *m/z*. Following loading of precursor lists and adjustment of method parameters, the flexImaging sequence from the MALDI-TIMS-$MS^1$ dataset was saved as a new imaging run file, and the laser offset was set to X=100, Y=0 (μm) in flexImaging, offsetting by the size of the laser. This offset allowed for the collection of TIMS-$MS^2$ spectra from unablated sample material while maintaining identical spatial distribution to the MALDI-TIMS-$MS^1$ dataset.

#### 5) Generation and Export of $MS^2$ Spectra

iprm-PASEF acquisition produces the same flexImaging (.mis) file as previously mentioned. Import into SCiLS Lab followed the same procedure and "Feature finding" parameters outlined in the '"Feature finding" and construction of feature lists' section. Upon opening the global feature table, the TIMS heatmap visualization will showcase all selected precursors as a jagged box, with fragment features denoted as small circles. The T-ReX[3] algorithm automatically designates any feature detected each respective defined precursor window as a 'precursor' ion in the global feature table. Any *m/z* detected outside of the defined precursor isolation window, but within the defined mobility isolation window, for each respective precursor is designated as a 'fragment' ion in the global feature list. Ions can be sorted based on the 'Isolation Window' column of the global feature table, grouping features based on their combined *m/z* and mobility windows. Features containing the same 'Isolation Window' were highlighted in the global feature table and used to generate separate feature lists for each TIMS-resolved $MS^2$ precursor/fragment list using the 'Create new feature list from selected rows' option in the feature table window. Despite the additional filtration by the mobility dimension, some *m/z* values labeled as 'fragment' ions in each of the precursor/fragment feature lists may be matrix or media contaminants. To rid the precursor/fragment feature lists of these contaminant features, the 'Find Values Co-Localized to Feature' tool in SCiLS Lab was used to filter out undesired features whose spatial distribution did not match that of the respective precursor (Table S7). Following matrix/media feature filtering, the generated filtered precursor/fragment feature lists were individually converted to MASCOT generic format (*.mgf) files using Python in combination with the SCiLS Lab API for downstream analysis and annotation of $MS^2$ spectra. The prototype tool can be found at: https://github.com/gtluu/SCiLS_Lab_iprm-PASEF_Exporter.

**Table S7.** “Find Values Co-Localized to Feature” parameters

| **Data** | |
|---|---|
| **Normalization** | Root Mean Square |
| **Use images of** | • Co_1-Co_4 = Co-Culture<br>• Gluta = *G. arilaitensis*<br>• P_solitum = *P. solitum*<br>• CCA = Cheese Curd Agar |
| **Correlate with feature** | 655.2727 m/z ± 20 ppm |
| **Feature list** | *Choose precursor/fragment feature list |
| **Save As** | *Specify name for filtered feature list |

*These parameters will depend on what feature lists are present and named within the SCiLS Lab dataset.

### 6) $MS^2$ Spectral Annotation

Following completion of all data acquisition steps, the generated *.mgf $MS^2$ spectra can be used to assign molecular annotations to the MALDI-TIMS-$MS^1$ dataset. The MALDI-TIMS-$MS^1$ MSI dataset was loaded into Bruker's MetaboScape software (version 2024b). A new feature table was created under the 'Project Manager' tab using the T-ReX[3] algorithm for feature finding. The previously generated Regions.srd file was loaded to provide MetabScape with the relevant spatial information from the MALDI-TIMS-$MS^1$ dataset. Region of Interest (RoI) subsampling was performed with the "Speckle size" set to 2x2, the "Max Speckles / RoI" set to 575, and the "Intensity Threshold" set to 500. Ion deconvolution was performed using the "T-Rex Default Single Spectra Positive Ions" method, monitoring for $[M+H]^+$, $[M+Na]^+$, $[M+K]^+$, $[M+Fe]^+$, $[M+Zn]^+$. No annotation methods were used upon generation of the initial feature table. After construction of the feature table, the generated *.mgf $MS^2$ spectrum file for each of the 12 selected precursors was uploaded to the respective feature in the table. "Spectral Library Annotation" was performed only for the features assigned $MS^2$ spectra.

**Table S8. GNPS2 Library Search Parameters**

| Name | Value |
|---|---|
| analog_search | 1 |
| blink_ionization | positive |
| blink_minpredict | 0.0075 |
| create_time | 2025-03-25 15:35:30 PDT-0700 |
| filtertostructures | 0 |
| fragment_tolerance | 0.1 |
| inputlibraries | LIBRARYLOCATION/LC/LIBRARY |
| inputspectra | USERUPLOAD/rashephe/Molecular Networking/250323/iPRM_molecular_networking/mgf_files |
| library_min_matched_peaks | 5 |
| library_min_similarity | 0.6 |
| peak_transformation | sqrt |
| pm_tolerance | 0.1 |
| search_algorithm | cos |
| searchtool | gnps |
| task | 6e2e3794132248aeb2dc6458ba67ea69 |
| topk | 20 |
| unmatched_penalty_factor | 0.6 |
| workflow_version | SERVER:2025.02.12;WORKFLOW:2025.02.03 |
| workflowname | librarysearch_workflow |

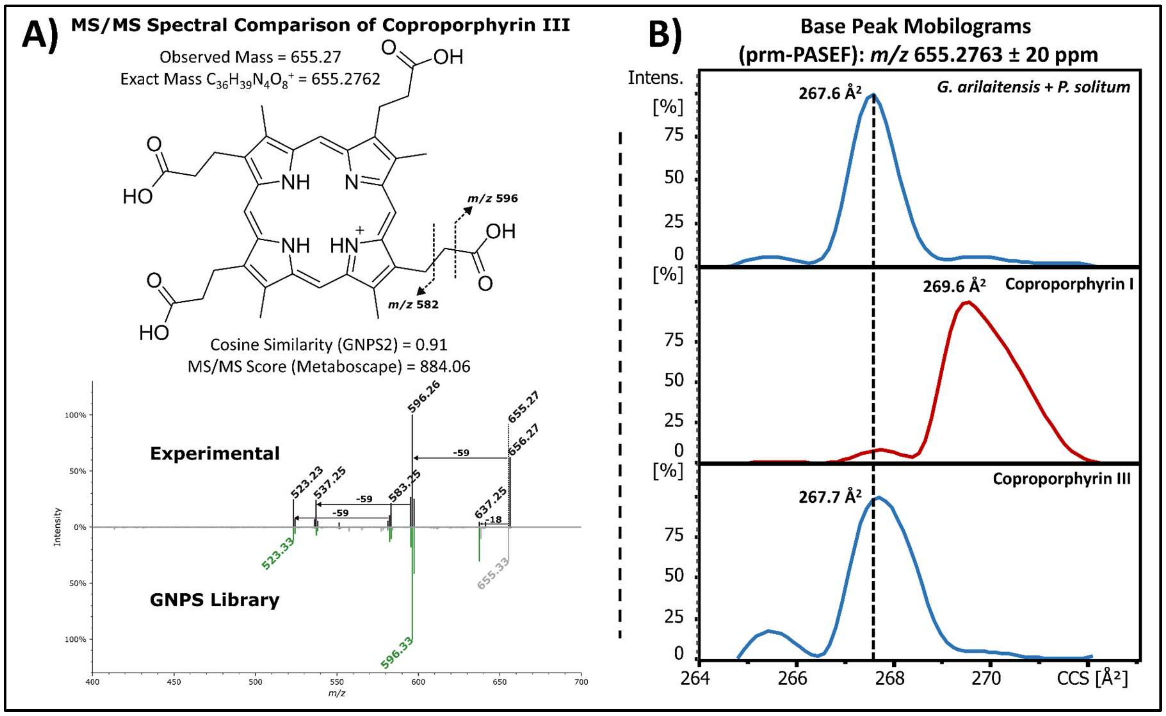


**Figure S1** Confirmation of coproporphyrin III in the *G. arilaitensis* and *Penicillium solitum* co-culture by $MS^2$ spectral annotation and TIMS resolution. A) $MS^2$ spectral annotation of coproporphyrin III. $MS^2$ fragmentation identified a coproporphyrin molecule, but cannot differentiate between coproporphyrin isomers. Previous work required lengthy HPLC co-injection analysis with standards to confirm the identity of the coproporphyrin isomer. B) Base Peak Mobilograms (BPMs) generated from MALDI iprm-PASEF spectra at *m/z* 655.2763 ± 20 ppm. The mobilogram peak from the co-culture region (top) matched well with the measured CCS of the major mobilogram peak of the coproporphyrin III standard (bottom). The major BPM peak for coproporphyrin I has minimal overlap with the associated CCS of coproporphyrin III, making them distinguishable by TIMS, albeit with a long ramp time (500 ms) and narrow mobility range ($1/K_0$ 1.29-1.32 $Vs/cm^2$).

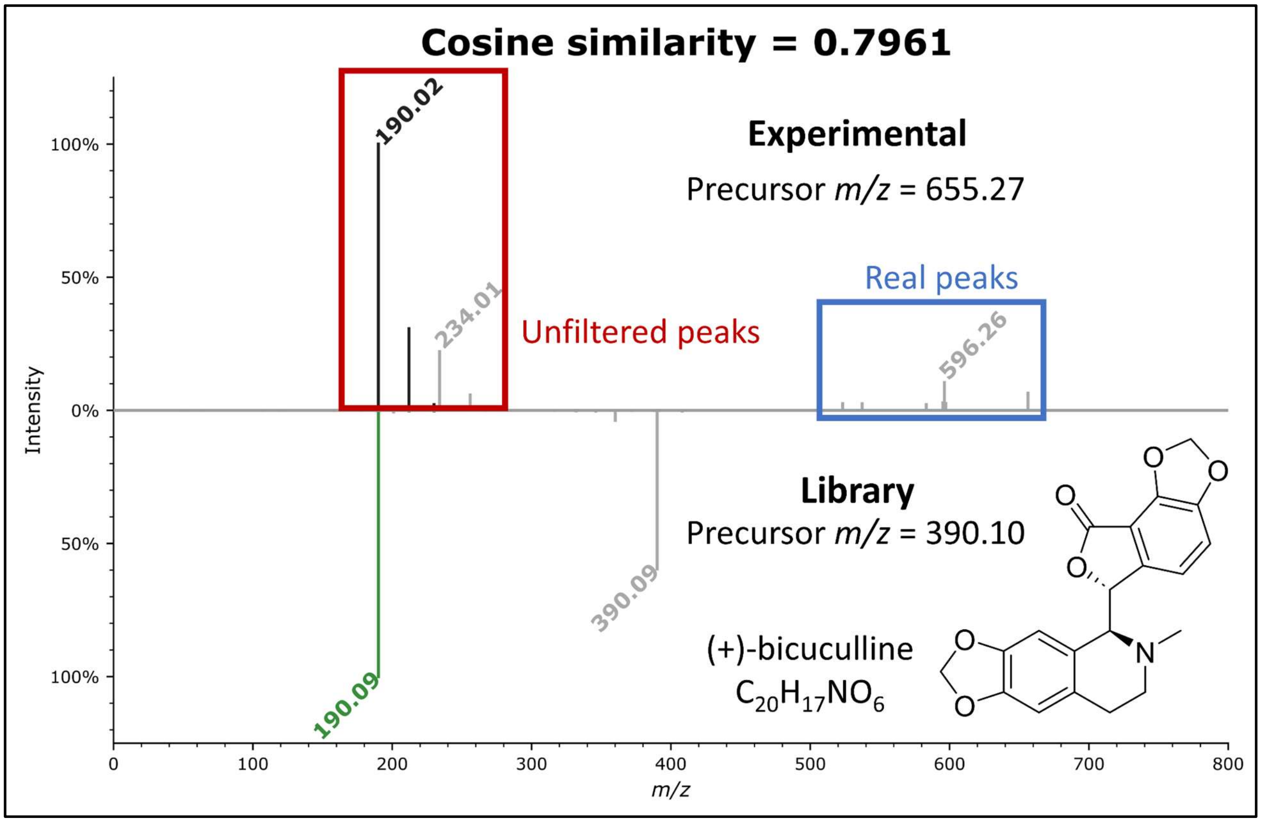


**Figure S2** Incorrect GNPS2 annotation of non-spatially filtered $MS^2$ spectrum for *m/z* 655.27. Recent software updates in SCiLS Lab have enabled MALDI iprm-PASEF data to be viewed, annotated and processed within the software. Prototypic versions of the MALDI iprm-PASEF workflow extracted all features within the same mobility window as the designated precursor without considering spatial information. As seen above, if there are non-precursor associated fragments within the same mobility window as ‘real’ fragment features, $MS^2$ library annotation results can be drastically skewed, decreasing annotation accuracy. Removal of the peaks highlighted in red through feature co-localization in SCiLS Lab prior to export of the *.mgf $MS^2$ spectrum yielded high-confidence spectral annotations for coproporphyrin as seen in Figure S1.

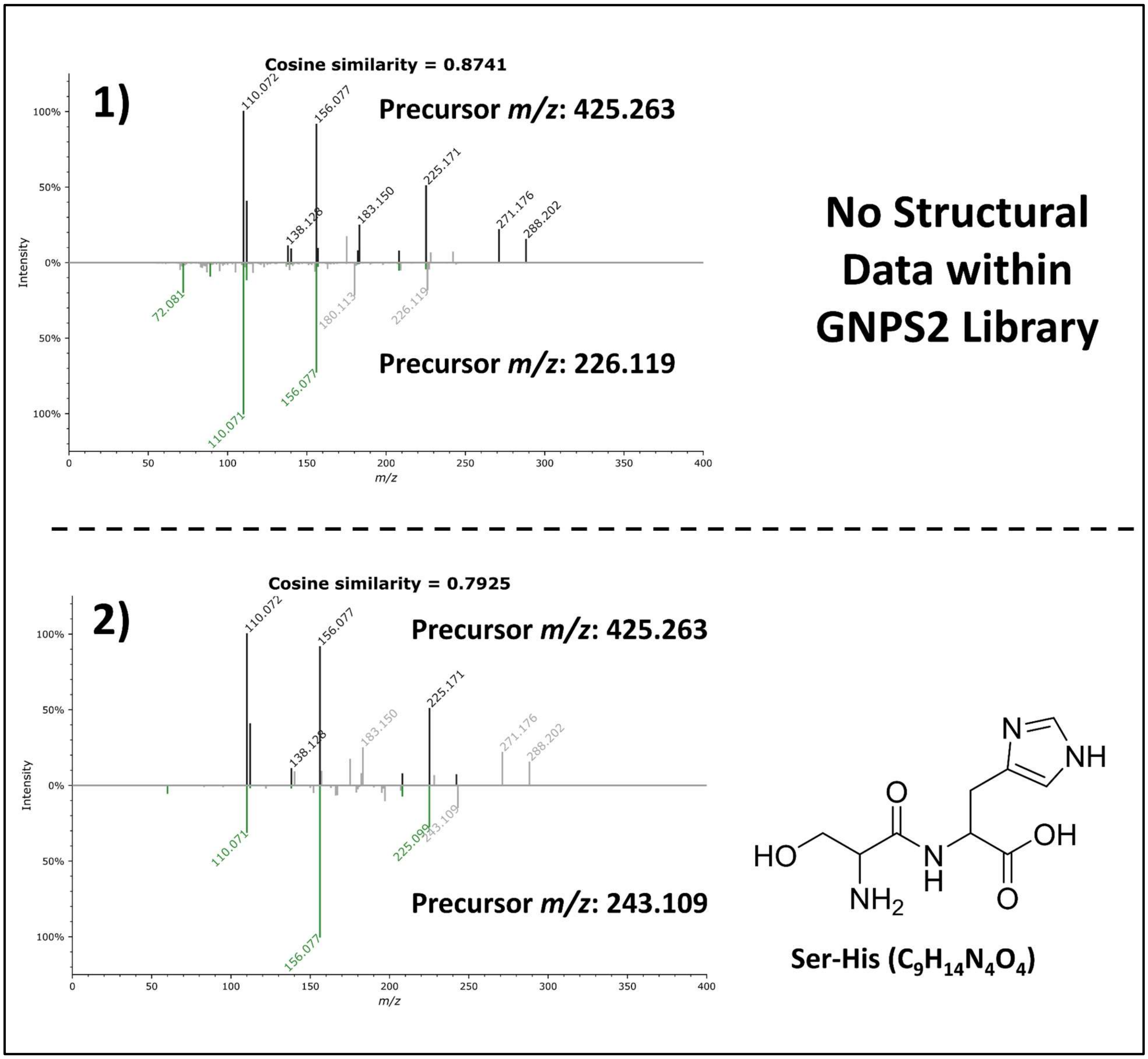


**Figure S3** Top GNPS2 library annotations for *m/z* 425.26. The top hit in the library did not have any structural data available for comparison, but the second highest library hit was for Ser-His, presumably due to matches with characteristic histidine fragments at *m/z* 156, which corresponds to protonated histidine, and likely means that the compound represented by this $MS^2$ spectrum has a C-terminal histidine that is being lost as the y1 fragment ion, consistent with Ser-His. The fragment at *m/z* 110 is also a characteristic histidine fragment, corresponding to the loss of CO and $H_2O$.[1] Due to the clear difference in precursor *m/z*, both GNPS2 library hits were deemed to be likely related, but not identical to the compound represented by *m/z* 425.26.

**Table S9. MetaboScape Spectral Library Annotation Method Parameters**

| **Spectral Library Annotation Method** | | | |
|---|---|---|---|
| **Search Libraries** | ALL_GNPS.msp | | |
| **Search Mode** | Parallel search | | |
| **Annotation on** | Complete Feature Table | | |
| **Settings** | | | |
| • Only allow annotations validated by MS/MS spectra<br>• Filter precursor m/z | | | |
| **Tolerances and scorings** | | | |
| | **Narrow** | **Wide** | **Unit** |
| *m/z* | 5 | 50.0 | ppm |
| mSigma | 20 | 100 | - |
| MS/MS score | 900 | 600 | - |
| CCS | 1.0 | 3.0 | % |

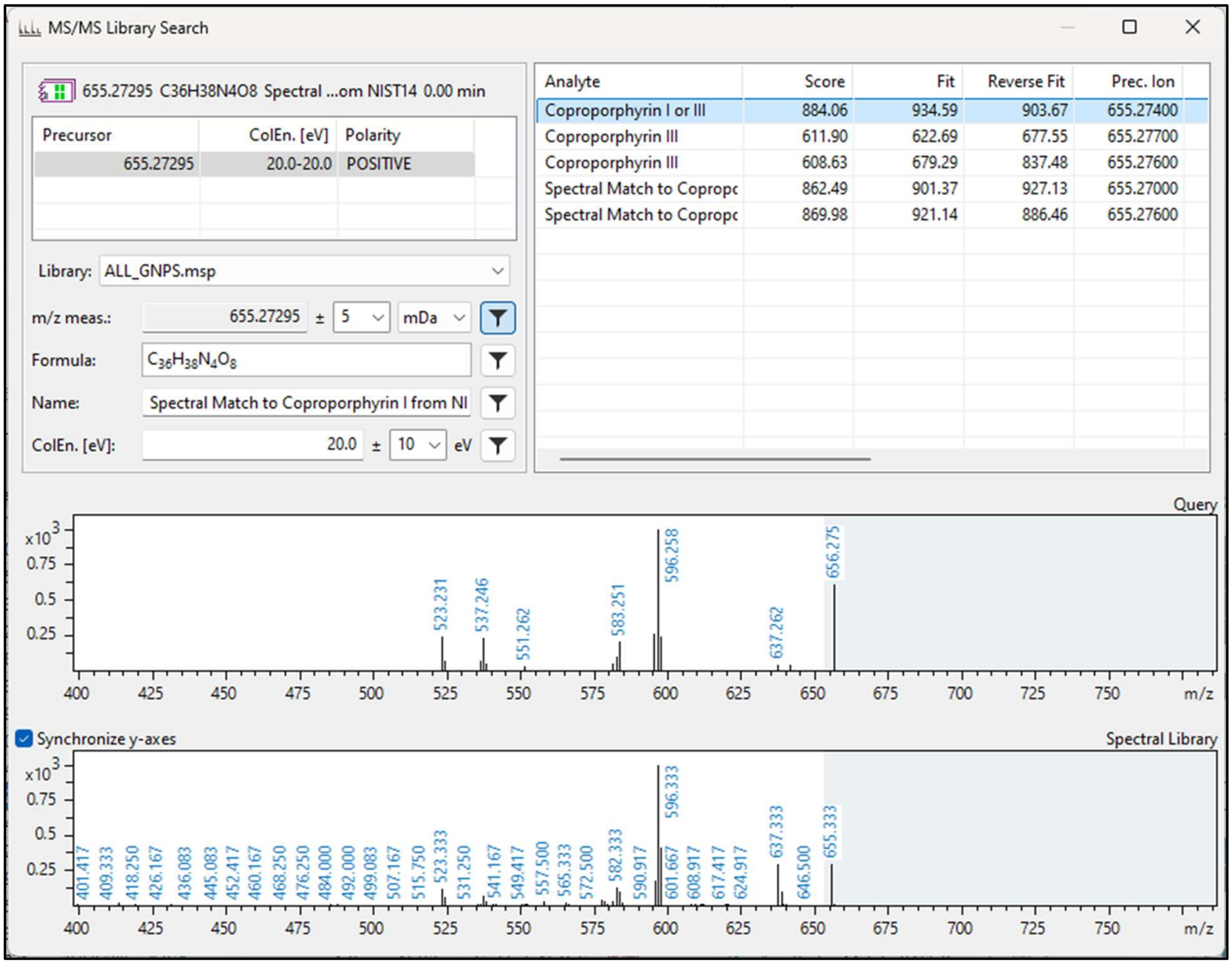


**Figure S4** MetaboScape spectral library search results for *m/z* 655.273. The top result from the $MS^2$ library search corresponded to 'Coproporphyrin I or III' with an MS/MS score of 884.06. This was consistent with the annotation we made in GNPS2 (Figure S1). Since coproporphyrin isomers showcase identical $MS^2$ fragmentation, annotation of the correct isomer is not possible with $MS^2$ spectra alone, and requires orthogonal validation (in this case via TIMS).

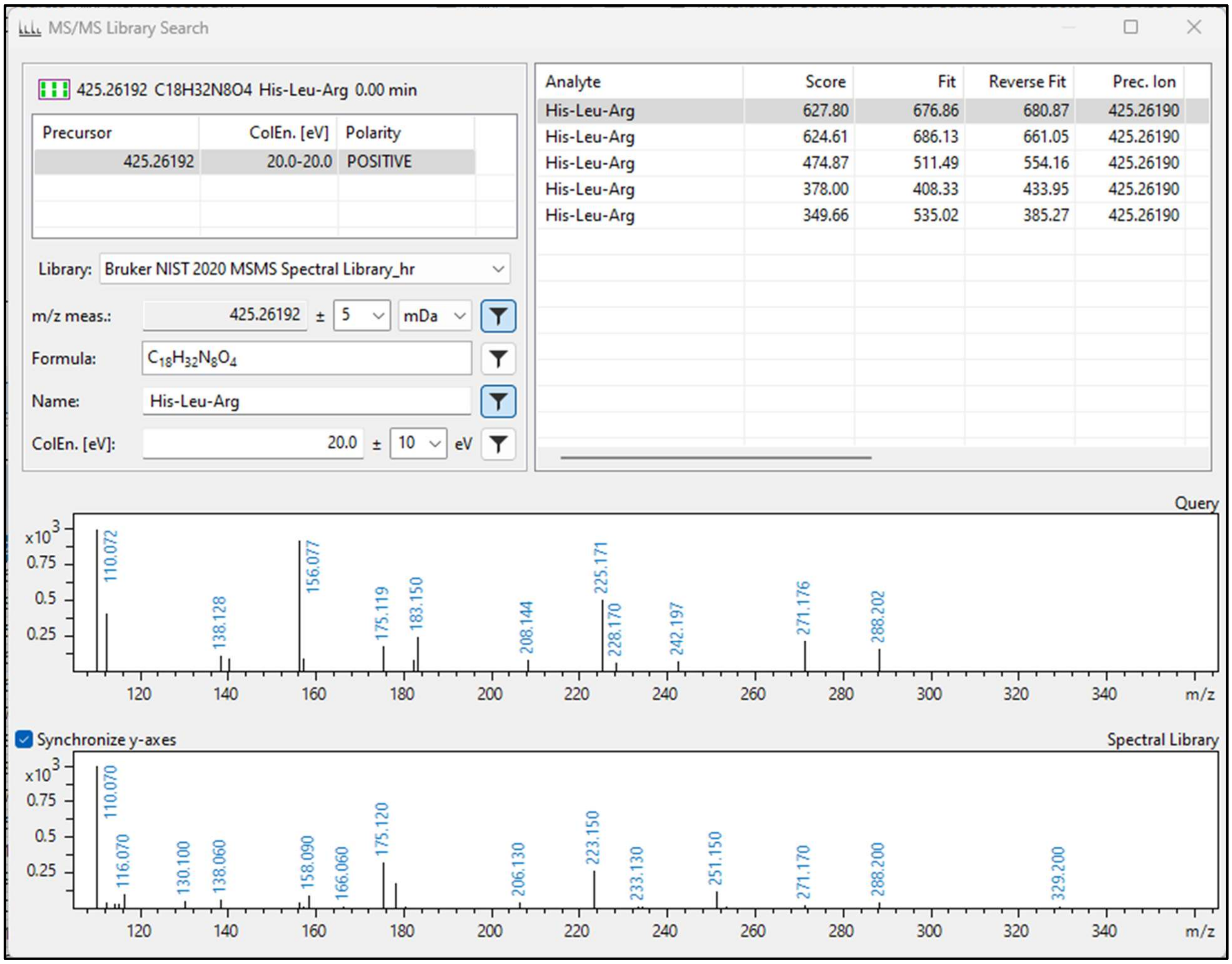


**Figure S5** MetaboScape Spectral Library annotation results for *m/z* 425.262. The top result was His-Leu-Arg from the Bruker NIST 2020 MS/MS spectral library with a relatively low-confidence MS/MS score of 627.80. Consistent with the GNPS2 data, the annotation remains of a peptidic nature, and contains a histidine residue. Additionally, the calculated intact mass of His-Leu-Arg ($[M+H]^+$=425.2619) is consistent with the observed precursor at *m/z* 425.262. Manual analysis of the $MS^2$ spectra however, would suggest a C-terminal rather than N-terminal histidine residue, as suggested by this annotation.

**Table S10. SIRIUS Compute Parameters**

<table>
<tr><th colspan="3">SIRIUS- Molecular Formula Identification</th></tr>
<tr><td colspan="3">General</td></tr>
<tr><td>Instrument</td><td colspan="2">Q-TOF</td></tr>
<tr><td>Filter by isotope pattern</td><td colspan="2">Yes</td></tr>
<tr><td>MS2 mass accuracy (ppm)</td><td colspan="2">10</td></tr>
<tr><td>MSMS isotope scorer</td><td colspan="2">IGNORE</td></tr>
<tr><td>Candidates stored</td><td colspan="2">10</td></tr>
<tr><td>Min candidates per ion stored</td><td colspan="2">1</td></tr>
<tr><td>Use DB formulas only</td><td colspan="2">all</td></tr>
<tr><td>Possible Ionizations</td><td colspan="2">$[M+H]^+$</td></tr>
<tr><td colspan="3">ILP</td></tr>
<tr><td>Tree timeout</td><td colspan="2">0</td></tr>
<tr><td>Compound timeout</td><td colspan="2">0</td></tr>
<tr><td>Use heuristic above m/z</td><td colspan="2">300</td></tr>
<tr><td>Use heuristic only above m/z</td><td colspan="2">650</td></tr>
<tr><td colspan="3">Elements allowed in Molecular Formula</td></tr>
<tr><td>Elements</td><td>Min # of atoms</td><td>Max # of atoms</td></tr>
<tr><td>H, C, N, O, P, S</td><td>0</td><td>inf</td></tr>
<tr><th colspan="3">CSI:FingerID - Fingerprint Prediction</th></tr>
</table>

| | |
|---|---|
| **Fallback Adducts** | all |
| **CSI:FingerID - Structure Database Search** | |
| **Search DBs** | all |
| **CANOPUS - Compound Class Prediction** | selected |

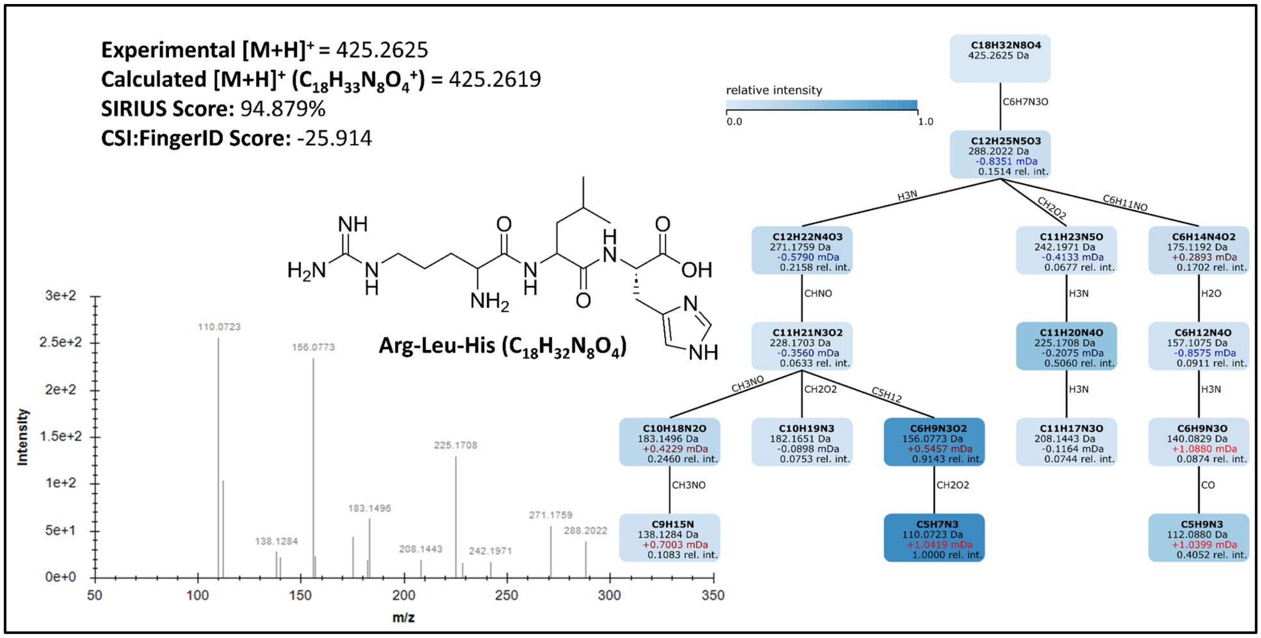


**Figure S6** SIRIUS Search results and MS/MS fragmentation tree. The top result in SIRIUS was Arg-Leu-His with a SIRIUS score of 94.879%. The calculated [M+H]+ of Arg-Leu-His is consistent with precursor *m/z* 425.2625. Lastly, this structural assignment is consistent with our hypothesis that *m/z* 425.2625 likely contains a C-terminal histidine residue based on the fragment at *m/z* 156, corresponding to a y1 histidine.

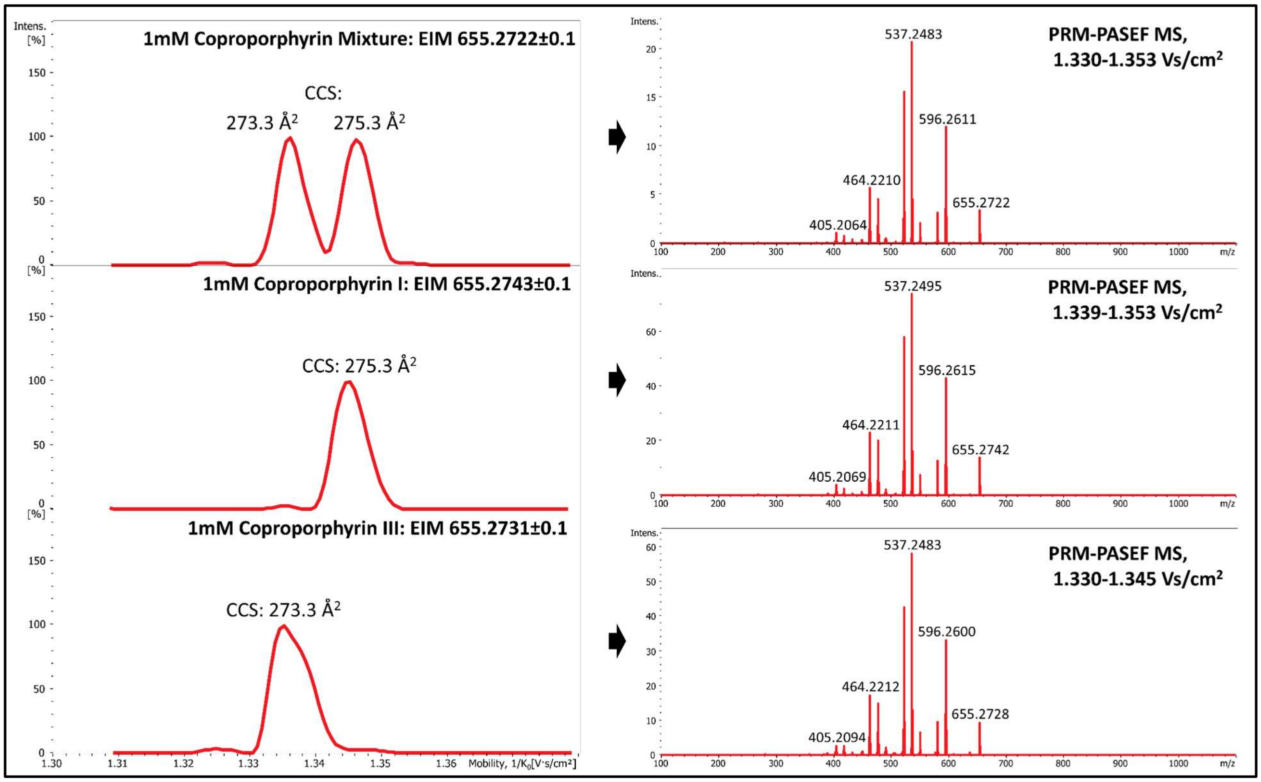


**Figure S7** Extracted ion mobilograms (EIMs) and averaged fragment spectra of *m/z* 655.27 from dried-droplet MALDI prm-PASEF data of coproporphyrin standards. When averaging the $MS^2$ fragmentation spectrum across both mobilogram peaks in the 1mM coproporphyrin mixture, the spectrum remains identical to the individual $MS^2$ spectra of coproporphyrin I and III. As can be observed above, coproporphyrin I and III can be resolved in the mobility dimension by extending the ramp time to 500 ms, and considerably narrowing the $1/K_0$ range, allowing for resolution of the two isomers. This however requires using an uncalibrated $1/K_0$ range that spans only 3 Vs/$cm^2$. Thus, $1/K_0$ values ,and therefore calculated CCS values, may experience some variation between MALDI-TIMS-MS acquisitions. For example, the CCS values for coproporphyrin I and III from our microbial MSI experiment (Figure 4 and S1) were 267.6 and 269.6 $Å^2$ respectively. Here, the values are shifted slightly, to 273.3 $Å^2$ for coproporphyrin I and 275.3 $Å^2$ for coproporphyrin III. That results in a 2.1% difference in experimental CCS between acquisitions. With such a minimal difference in mobility between coproporphyrin I and III, it might be necessary to include standards in future analyses to account for instrument drift.

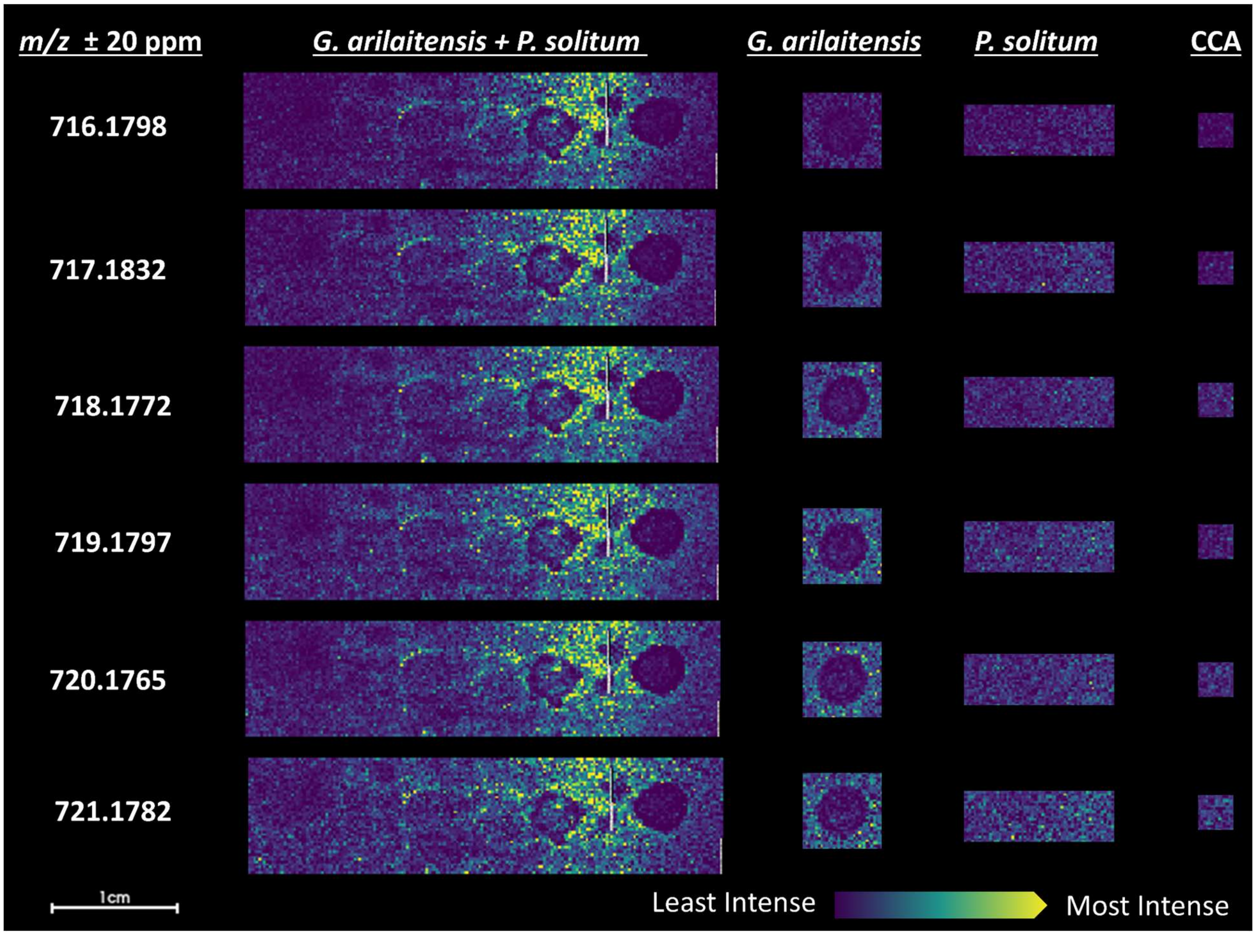


**Figure S8** MALDI-TIMS-MS[1] ion images of previously reported isotopologue *m/z*'s for zinc-coproporphyrin. These data are consistent with the spatial distribution and reported *m/z* values previously reported for zinc-coproporphyrin in co-culture by Cleary et al.[2], but no further validation was performed specifically for zinc-coproporphyrin analogs in this study. The monoisotopic mass of zinc-coproporphyrin ($C_{36}H_{36}N_4O_8Zn^+$) is 716.1825 (mass error = 3.77 ppm).

### References


(1) Zhang, P.; Chan, W.; Ang, I. L.; Wei, R.; Lam, M. M. T.; Lei, K. M. K.; Poon, T. C. W. Revisiting Fragmentation Reactions of Protonated α-Amino Acids by High-Resolution Electrospray Ionization Tandem Mass Spectrometry with Collision-Induced Dissociation. *Sci. Rep.* **2019**, *9* (1), 6453.

(2) Cleary, J. L.; Kolachina, S.; Wolfe, B. E.; Sanchez, L. M. Coproporphyrin III Produced by the Bacterium *Glutamicibacter Arilaitensis* Binds Zinc and Is Upregulated by Fungi in Cheese Rinds. *mSystems* **2018**, *3* (4). https://doi.org/10.1128/msystems.00036-18.
